# Combinational adjuvants delivered by ink-jet potentiate naked mRNA vaccines for robust protection against infectious diseases

**DOI:** 10.64898/2025.12.24.696305

**Authors:** Nan Qiao, Junichi Ishikawa, Fumihiko Yasui, Kaori Sano, Kei Miyakawa, Akimasa Hayashi, Ahmad Faisal Amiry, Liu Haonan, Zhu Xueyang, Taku Sato, Renata Pasqualini, Wadih Arap, Kazunori Kataoka, Hideki Hasegawa, Michinori Kohara, Satoshi Uchida

## Abstract

The use of lipid nanoparticles (LNPs) to enhance delivery and immunogenicity of mRNA vaccines raises safety concerns due to strong reactogenicity. Developing naked mRNA vaccines, which can be delivered through liquid jet-injection to the skin, may offer a safer alternative, but their immunogenicity has been limited. To address this challenge, we screened clinically relevant adjuvants to enhance the potency of naked mRNA jet-injection vaccines in mouse models. Although many adjuvants showed little or inhibitory effects on inducing humoral or cellular immunity—despite their known ability to potentiate conventional non-mRNA vaccine platforms—we identified a combination of aluminum phosphate and CpG oligonucleotide that substantially enhanced both humoral and cellular immune responses with increased IgG2a/IgG1 ratio. This adjuvanted naked mRNA vaccine induced IgG2a and cellular immunity at levels comparable to those achieved with LNPs but with lower local and systemic reactogenicity. Mechanistic investigations revealed a key role for type I interferon signaling during immunization. Notably, this vaccine formulation in mice virtually eliminated infection by both SARS-CoV-2 and influenza A virus (A/California/7/2009 H1N1) in plaque-forming assays. Collectively, the adjuvanted naked mRNA jet-injection vaccine represents a novel, effective, and safe platform against infectious diseases, with potential for clinic-ready translational applications.

## Introduction

mRNA vaccines against COVID-19 have demonstrated high efficacy and acceptable safety profiles in both clinical trials and real-world settings (1–4), establishing their utility as a platform for infectious disease prevention. Their application is expanding to other viruses and bacteria (5–8), with promising outcomes reported in several clinical trials (9, 10). However, the reactogenicity of mRNA vaccines, such as local pain and fever, tends to be stronger and more frequent than that of conventional vaccines (11). Moreover, recent clinical trials have also revealed an extremely high incidence of chronic urticaria (10, 12). Finally, although rare, life-threatening reactions such as anaphylaxis and myocarditis have been reported (13, 14). Although these issues may be acceptable for highly life-threatening pathogens (as defined by the World Health Organization and/or the Centers for Disease Control and Prevention), they must be addressed to enable broader application of mRNA vaccines across various infectious diseases and to reduce vaccine hesitancy (15).

Lipid nanoparticles (LNPs), which encapsulate the mRNA and enhance delivery and immunogenicity, are a key contributor to reactogenicity (16–18), although the mechanisms underlying these adverse reactions remain incompletely understood. Both animal and human studies have shown that LNPs induce strong proinflammatory responses in vivo (16, 19, 20). Moreover, antibody responses to LNP components may contribute to anaphylaxis and chronic urticaria (21, 22). The biodistribution of LNPs is also uncontrolled, with locally injected LNPs leaking into systemic circulation and migrating to distant organs (23–25). Clinical data suggest a potential link between systemic LNP distribution and severe adverse events, including myocarditis and autoimmune hepatitis (26, 27). These concerns have prompted efforts to develop less inflammatory LNPs with controlled biodistribution (28–30), although fully resolving these issues remains a major challenge.

The development of naked mRNA-based vaccines presents a straightforward alternative strategy to overcome the safety concerns associated with LNPs. However, this approach requires techniques to enhance the efficacy of naked mRNA vaccines to levels comparable to LNP-based formulations by incorporating LNP-like functionalities. One of the key roles of LNPs in mRNA vaccines is the efficient delivery of mRNA into antigen-presenting cells (APCs) (18, 31, 32). To replicate this delivery efficiency with naked mRNA, we previously employed jet-injection targeting the skin (33), which is rich in APCs. This method markedly enhanced the immunogenic potential of naked mRNA vaccines against infectious diseases in both mice and non-human primates. Of note, this injector has already been used in clinical trials for plasmid DNA vaccines and caused only limited local reactions (34), demonstrating strong potential for translation into clinical applications. Nevertheless, despite the high mRNA delivery efficiency, antibody production induced by naked mRNA vaccines remains lower than that achieved with LNPs, underscoring the need for additional factors to enhance vaccine efficacy.

The immunostimulatory properties of LNPs, which act as built-in adjuvants, are another critical factor contributing to their high vaccination efficacy (17–20, 35). Therefore, in this study, we combined jet-injection technology with immunostimulatory adjuvants and conducted a systematic screening to identify adjuvants that enhance the efficacy of naked mRNA vaccines. We focused on licensed adjuvants and their derivatives to minimize safety concerns and facilitate the clinical translation. Interestingly, many adjuvants that enhance the efficacy of conventional inactivated, toxoid, or recombinant protein vaccines induced little or even inhibitory effects on mRNA vaccine performance, highlighting the complexity of adjuvant selection for mRNA platforms. Nevertheless, our screening identified an optimal combination of adjuvants that increased antibody production by 10- to 100-fold, elevated the IgG2a/IgG1 ratio and simultaneously enhanced cellular immune responses in mice. Notably, the levels of IgG2a and cellular immunity were comparable to those induced by LNPs, while adjuvanted naked mRNA elicited lower local and systemic reactogenicity. Mechanistic analyses revealed a critical role for type I interferon (IFN) signaling in the immunization process of adjuvanted naked mRNA vaccines. Ultimately, this system provided robust protection against SARS-CoV-2 and influenza virus (A/California/7/2009 H1N1), reducing viral loads to background levels in plaque-forming assays, again demonstrating its potential for combating infectious diseases with well-documented epidemic or pandemic potential.

## Results

### Screening reveals divergent effects of adjuvants on mRNA vaccine performance

In selecting adjuvants, we focused on licensed compounds or their derivatives, including aluminum phosphate (Al-Phos), MF59-like adjuvant (MF59-l), AS03-like adjuvant (AS03-l), lipopolysaccharide (LPS), and CpG oligodeoxynucleotides (CpG) (36–38). Aluminum-based adjuvants have a long precedent history of use in human vaccines, demonstrating effectiveness in toxoid, inactivated, and recombinant protein vaccines. Among their derivatives, we selected negatively charged Al-Phos to avoid electrostatic interactions with the negatively charged mRNA (39). MF59 and AS03 are licensed for use in inactivated influenza vaccines and are composed of oil-in-water emulsions. MF59 contains squalene, while AS03 includes both squalene and α-tocopherol as immunostimulants. LPS is a Toll-like receptor 4 (TLR4) agonist used in AS01 and AS04, which are licensed for recombinant protein vaccines. CpG stimulate TLR9 and are licensed for recombinant protein-based hepatitis B vaccines.

The addition of these adjuvants to naked mRNA vaccines could influence the delivery process via jet-injection through several mechanisms. Physical interactions with mRNA may impair delivery, while innate immune activation could reduce protein translation efficiency (30, 40, 41). To assess these possibilities, we first evaluated the impact of adjuvants on protein expression by using *luciferase* mRNA as a reporter. For injection, we used a needle-free liquid jet injector powered by a little gunpowder, which safely and efficiently delivers mRNA into the skin, a tissue rich in APCs (42). Following jet-injection into mice, most adjuvants had minimal impact on luciferase expression, except MF59-l, which reduced expression levels by 2- to 5-fold (**Figure 1A, B**). Notably, efficient protein expression persisted for at least three days regardless of adjuvant addition.

**Figure 1.**
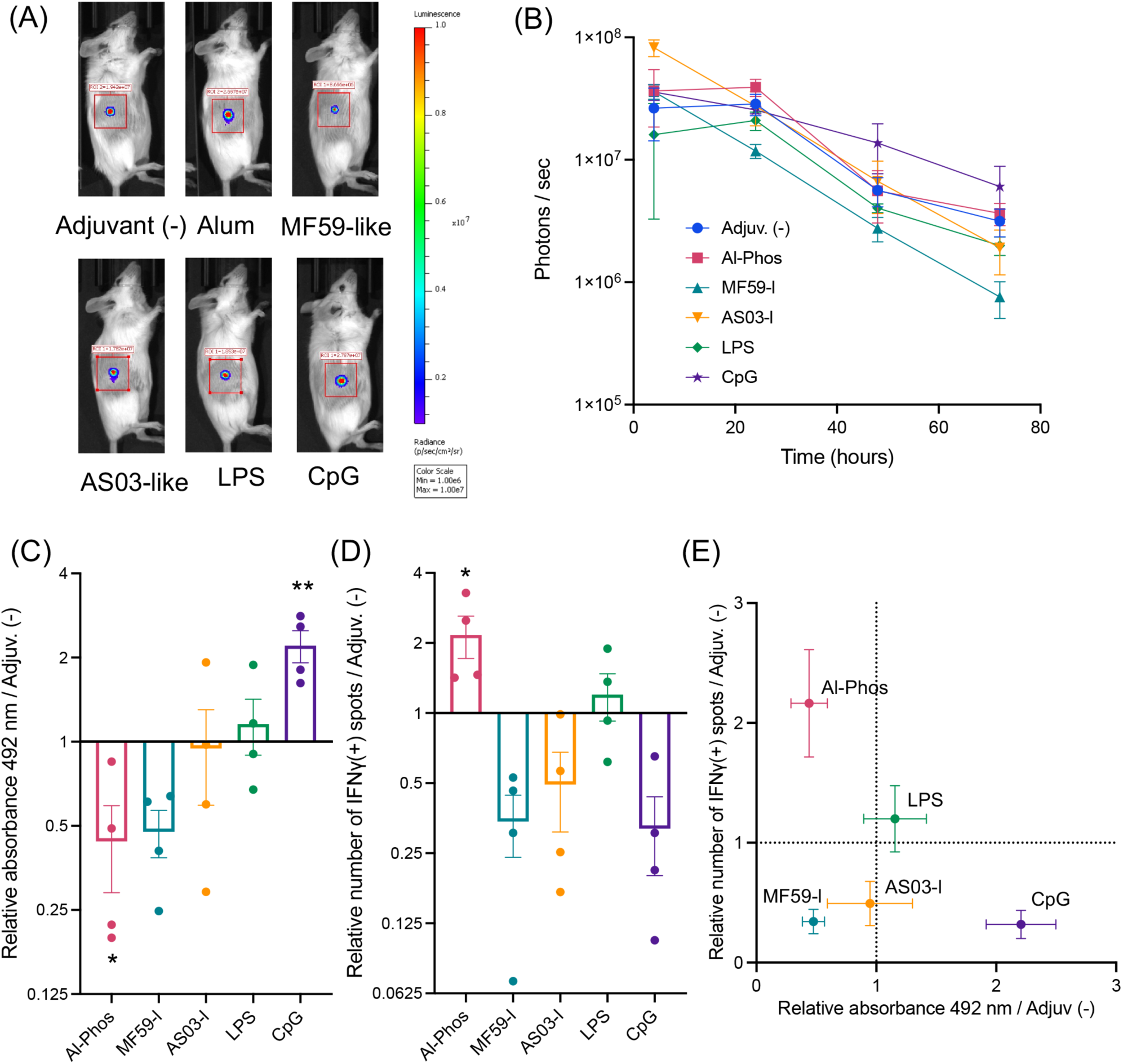
Influence of adjuvants on mRNA delivery and vaccination efficiency in naked mRNA jet-injection. (**A, B**) Luciferase expression efficiency. Mice received jet-injection of naked *luciferase* mRNA alone or with different adjuvants. (**A**) Representative in vivo imaging system (IVIS) images of luciferase expression 24 hours post-injection. (**B**) Luciferase expression levels at the injection site quantified by IVIS imaging at 4, 24, 48, and 72 hours-post-injection. *n* = 4. (**C–E**) Vaccination efficacy. Mice were immunized with naked *spike* mRNA with or without adjuvants via jet-injection, twice at a 3-week interval. Efficacy was evaluated two weeks after the second dose. (**C**) Fold change in spike-specific total IgG levels in plasma relative to the group without adjuvant. *n* = 4. (**D**) Fold change in IFN-γ-producing splenocytes by ELISpot relative to the group without adjuvant. *n* = 4. (**E**) Correlation between IFN-γ–producing splenocytes and spike-specific total IgG levels. *n* = 4. Data are presented as mean ± standard error of the mean (s.e.m.) Raw data for (**C-E**) are shown in **Supplementary Figure S1**. *: *p* < 0.05; **: *p* < 0.01 versus the group without adjuvant. See **Supplemental Figure S1** for details of statistical analyses.

Next, we evaluated the immunological effects of these adjuvants in a vaccine targeting the SARS-CoV-2 spike protein. Mice were vaccinated twice at a three-week interval, and anti-spike antibody levels in plasma were measured by using ELISA, while cellular immune responses were assessed via ELISpot of splenocytes. To visualize the impact of adjuvant addition, results are presented as relative values compared to the group treated with *spike* mRNA alone without adjuvants. Specifically, **Figure 1C** shows humoral immune responses as relative ELISA absorbance values, and **Figure 1D** shows cellular immune responses as relative numbers of IFN-γ–secreting splenocytes in ELISpot. The raw data used to generate these figures are provided in **Supplemental Figure S1**. Many adjuvants showed little or even inhibitory effects on vaccine efficacy. Specifically, Al-Phos and MF59-l decreased antibody production, while MF59-l, AS03-l, and CpG reduced cellular immune responses. Furthermore, the effects of each adjuvant on humoral and cellular immunity were poorly correlated (**Figure 1E**), highlighting the complexity of adjuvant function in mRNA vaccines. Notably, CpG showed a positive effect on antibody production, and Al-Phos enhanced cellular immune responses. Based on these findings, we next tested the combination of Al-Phos and CpG.

### Combinational adjuvants enhance both humoral and cellular immune responses

To evaluate the efficacy of combinational adjuvants, we administered an admixture of naked *spike* mRNA, Al-Phos, and CpG via jet-injection into mice, with two doses given three weeks apart. Five weeks after the initial vaccination, the combination of Al-Phos and CpG significantly enhanced both humoral and cellular immune responses, as evidenced by elevated anti-spike IgG levels in ELISA and increased numbers of spike-reactive splenocytes in ELISpot assays [*p* < 0.0001 for total IgG and IgG2a; *p* =0.0137 for IgG1; and *p* = 0.0126 in ELISpot compared with Adjuvant (-); one-way analysis of variance (ANOVA) followed by Tukey’s tests; **Figure 2A–D**]. This enhancement contrasts sharply with the use of individual adjuvants, which failed to simultaneously improve both humoral and cellular responses (**Figure 1C–E**). Notably, Al-Phos alone inhibited antibody production, while CpG alone impaired cellular immunity. In contrast, their combination effectively boosted both arms of the immune response.

**Figure 2.**
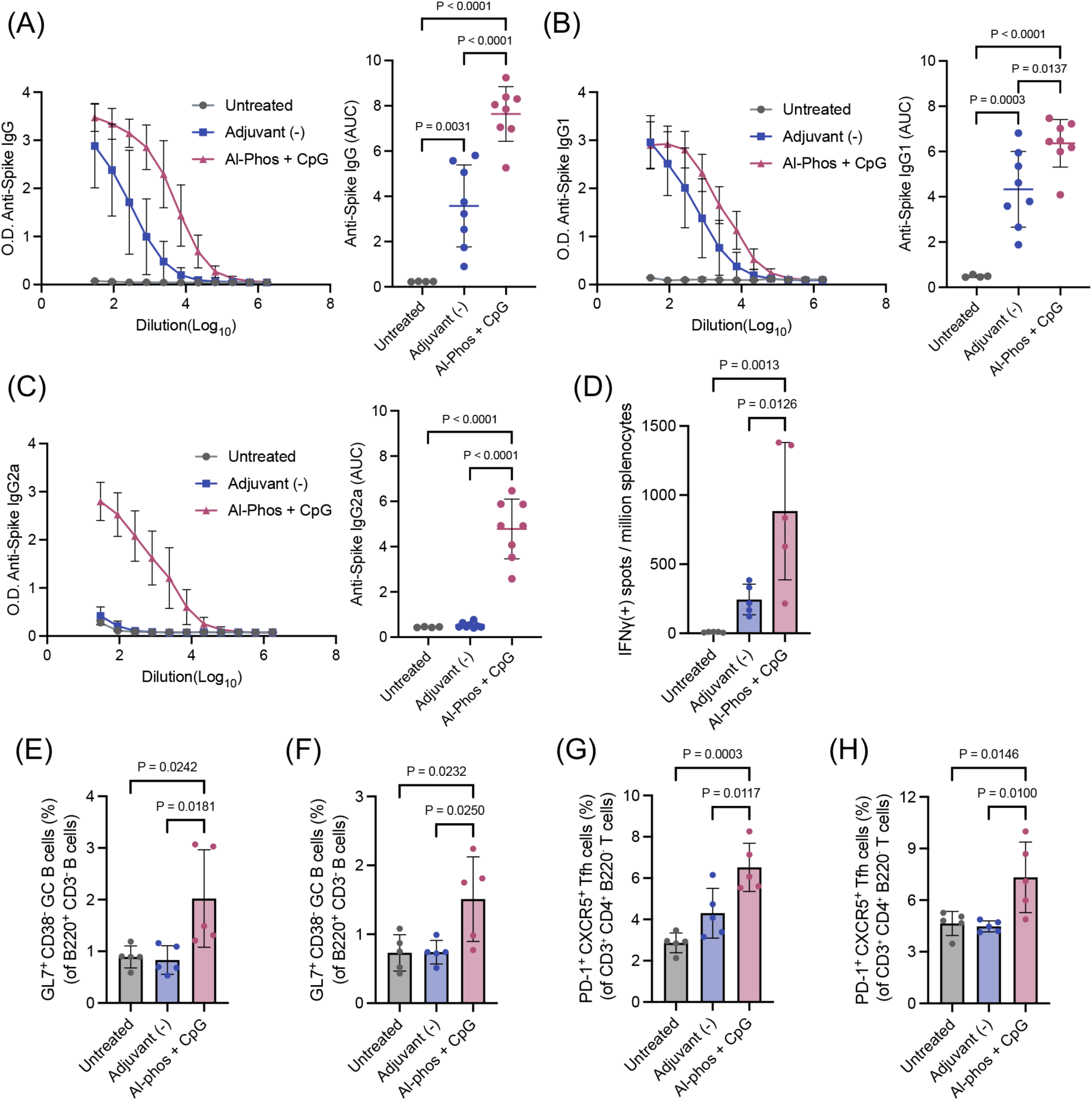
Immunization efficiency of naked mRNA jet-injection with combinational adjuvants. (**A–D**) Vaccination efficacy. Mice were immunized with naked *spike* mRNA with or without Al-Phos and CpG adjuvants via jet-injection, twice at a 3-week interval. Blood and spleens were collected 2 weeks after the second dose. (**A–C**) Antibody responses in plasma: (**A**) IgG, (**B**) IgG1, and (**C**) IgG2a. Left: Optical density (O.D.) from ELISA of serially diluted plasma. Right: Area under the curve (AUC) of the corresponding dilution curves. (**D**) IFN-γ–secreting splenocytes quantified by ELISpot. *n* = 4 for untreated, *n* = 8 for Adjuvant (-) and Al-Phos + CpG groups. (**E–H**) Germinal center formation in dLNs. dLNs were collected 7 days after jet-injection. (**E, F**) Percentage of germinal center (GC) B cells in inguinal LN (**E**) and axillary LN (**F**). *n* = 5. (**G, H**) Percentage of T follicular helper (Tfh) cells in inguinal LN (**G**) and axillary LN (**H**). *n* = 5. See **Supplemental Figure S2** for the gating strategy. Data are presented as mean ± standard deviation (s.d.), with statistical analyses performed by using one-way ANOVA followed by Tukey’s multiple comparisons test.

IgG subclass analysis revealed that the naked mRNA vaccine without adjuvants primarily induced IgG1, with only modest levels of IgG2a (**Figure 2B, C**). This low IgG2a/IgG1 ratio is indicative of a Th2-skewed immune response, which has been associated with an increased risk of vaccine-associated enhanced disease (VAED) in infectious disease vaccines (43, 44). In contrast, the combinational adjuvants induced a more pronounced increase in IgG2a relative to IgG1, resulting in higher IgG2a/IgG1 ratio while increasing levels of both subclasses. This balanced IgG2a/IgG1 response may help mitigate the risk of VAED.

To gain mechanistic insight into the immunological processes influenced by the combinational adjuvants, we evaluated the induction of T follicular helper (Tfh) cells and germinal center (GC) B cells in the draining lymph nodes (dLNs) at 7 days post-vaccination. The adjuvanted formulation significantly increased the proportions of both Tfh and GC B cells compared to untreated mice and those receiving unadjuvanted naked mRNA [*p* < 0.05 compared with Adjuvant (-); one-way ANOVA followed by Tukey’s tests; **Figure 2E–H**; **Supplemental Figure S2**]. In contrast, mRNA alone did not elevate Tfh or GC B cell levels. These findings indicate that the adjuvanted naked mRNA vaccine successfully induces GC responses, which are essential for immunoglobulin class switching and affinity maturation.

### Type I IFN responses play a critical role in vaccination with adjuvanted naked mRNA

To further elucidate the mechanisms underlying the effective adjuvanticity of the Al-Phos and CpG combination, we performed transcriptome analysis of the injection-site skin and dLNs 24 hours post-jet-injection of naked mRNA with or without adjuvants by using RNA sequencing. Gene Set Enrichment Analysis (GSEA) based on the Molecular Signatures Database (MSigDB) revealed enrichment of general and specific immune signaling pathways in the combinational adjuvant group compared to the group jet-injected with naked mRNA without adjuvants (**Figures 3A–D**). Specifically, pathways related to IFN-γ response, IFN-α response, TNF-α signaling via NF-κB, and IL-6/JAK/STAT3 signaling were enriched in the adjuvanted group in both skin and dLNs. Among these, previous reports have highlighted the critical roles of type I IFN and IL-6 pathways in potentiating mRNA vaccination through their adjuvanticity (20, 35, 45–47). Volcano plots and expression profiling of differentially expressed genes (DEGs) further revealed enhanced expression of genes related to these pathways, as well as other cytokines, IFNs, chemokines, their downstream signaling molecules, and inflammasome components in the adjuvanted group (**Figures 3E–H**). Particularly, numerous downstream molecules of type I IFN were upregulated by the combinational adjuvants.

**Figure 3.**
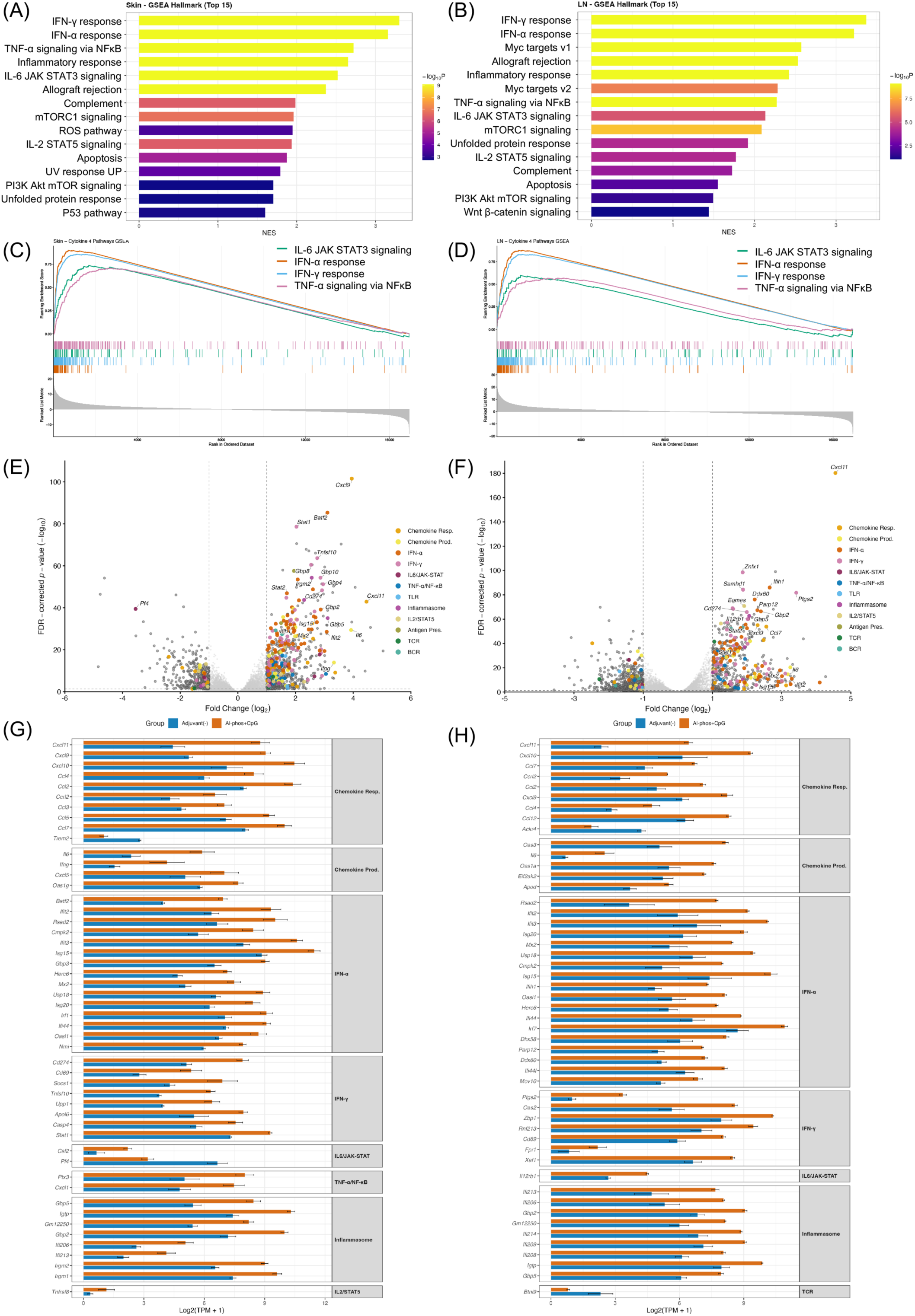
Transcriptomic analysis after adjuvanted naked mRNA vaccination. Mice received naked mRNA with or without Al-Phos and CpG adjuvants via jet-injection. At 24 hours post-immunization, injection site skin and dLN were collected for RNA sequencing. Transcriptomes were compared between groups with and without adjuvants. *n* = 3. (**A-D**) Gene Set Enrichment Analysis (GSEA) by using MSigDB Hallmark gene sets. (**A, B**) Top 15 enriched pathways ranked by normalized enrichment score (NES). (**C, D**) GSEA enrichment for each pathway. Top panels: Running enrichment scores across ranked genes. Middle panels: Positions of pathway genes. Bottom panels: Wald statistic distribution. (**A, C**) Skin. (**B, D**) dLN. (**E, F**) Volcano plots of differentially expressed genes (DEGs) in skin (**E**) and dLN (**F**). Genes with |log_2_FC| ≥ 1 and false-discovery rate (FDR) p < 0.05 are colored by 12 immune pathway categories derived from MSigDB Hallmark and GO:BP terms. (**G, H**) Expression profiles of pathway-associated DEGs (top 50 by |log_2_FC|) in skin (**G**) and dLN (**H**), grouped by pathway category.

Among these enriched pathways, we focused on type I IFN and IL-6 signaling for further mechanistic evaluation due to their reported contributions to the adjuvanticity of LNP-based mRNA vaccines (20, 35, 45–47). First, quantitative PCR analysis of cytokines was performed in the injected skin and dLNs to validate the RNA sequencing findings. The Al-Phos and CpG adjuvants enhanced *IFN-β* and *IL-6* transcript expression compared to vaccination without adjuvants 24 hours post-jet-injection of naked mRNA (**Figures 4A–C**), consistent with the RNA sequencing results. The extent of enhancement in lymph nodes was more pronounced for *IFN-β* than for *IL-6*. Notably, the combinational adjuvants did not increase *IL-1β* expression, a key molecule associated with mRNA vaccine adjuvanticity (48), in dLNs, although *IL-1β* expression was elevated in the skin.

**Figure 4.**
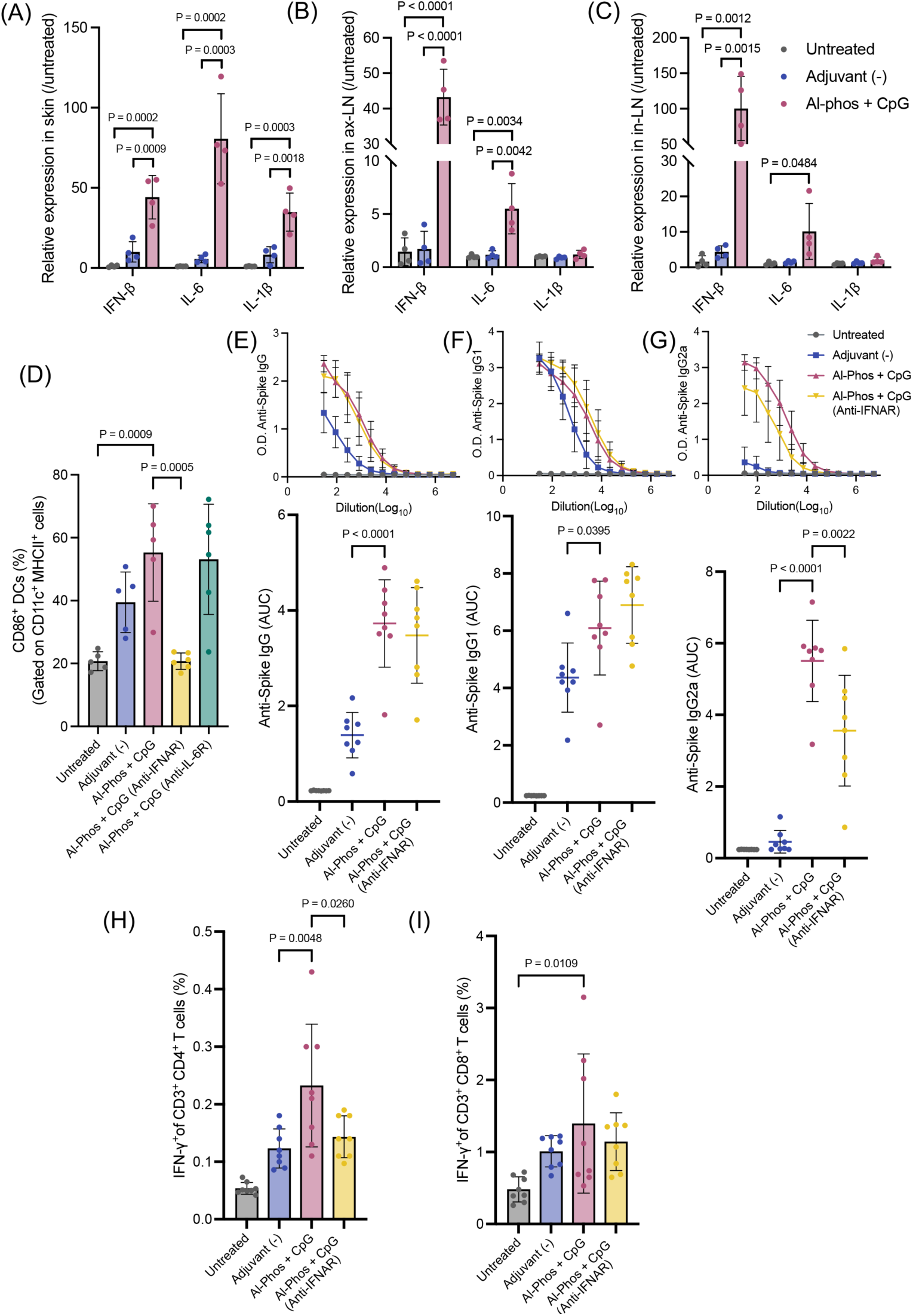
Signaling pathways involved in the vaccination process. (**A–C**) Cytokine expression in skin (**A**), axillary LN (ax-LN) (**B**), and inguinal LN (in-LN) (**C**). Transcript levels of *IL-6, IFN-β*, and *IL-1β* were measured 24 hours after naked mRNA jet-injection. *n* = 4. (**D**) Effect of blocking immune signaling on DC activation. Mice were intraperitoneally injected with anti-IFNAR or anti-IL-6R antibodies one day before receiving naked mRNA jet-injection with or without adjuvants. One day post-injection, flow cytometry of cells in dLNs was performed to measure CD86 expressed in CD11c^+^ DCs. *n* = 6. (**E-I**) Effect of blocking type I IFN signaling on vaccination efficacy. Mice were immunized with *spike* mRNA twice at a three-week interval, and immunological evaluations were performed at two weeks after the second vaccination. Anti-IFNAR antibodies were administered one day prior to each vaccination. (**E–G**) Antibody responses in plasma: (**E**) IgG, (**F**) IgG1, and (**G**) IgG2a. Top panels: Optical density (O.D.) from ELISA of serially diluted plasma. Bottom panels: Area under the curve (AUC) of the corresponding dilution curves. (**H, I**) T cell responses. (**H**) CD4^+^ T cells and (**I**) CD8^+^ T cells expressing IFN-γ after antigen stimulation were quantified by intracellular cytokine staining and flow cytometry. *n* = 8. See **Supplemental Figures S3 and S4** for the gating strategy. Data are presented as mean ± s.d. with statistical analyses performed by using one-way ANOVA followed by Tukey’s multiple comparisons test.

To investigate the contribution of type I IFN and IL-6 signaling further, we administered blocking antibodies against the IFN-α/β receptor (IFNAR) or IL-6 receptor one day prior to vaccination. At one day post-vaccination, we assessed dendritic cell (DC) activation, an essential step in immunization, by measuring the percentage of CD86⁺ cells among CD11c^⁺^ MHCII^⁺^ DCs in the dLNs. While jet-injection of naked *spike* mRNA modestly activated DCs even without adjuvants, the combinational adjuvants further increased CD86 expression demonstrating their capability to stimulate DCs (**Figure 4D, Supplemental Figure S3**). Notably, pre-injection of anti-IFNAR antibodies completely abrogated DC activation, reducing the percentage of CD86^⁺^ DCs to levels comparable to untreated controls. In contrast, pre-injection of anti-IL-6 antibodies had no detectable effect on DC activation.

We next examined the involvement of type I IFN signaling in adaptive immunity. Mice were vaccinated with *spike* mRNA twice at a three-week interval, with anti-IFNAR antibodies administered one day before both the prime and boost doses. In immunological analyses performed two weeks after the boost, anti-IFNAR treatment did not affect anti-spike total IgG or IgG1 levels but markedly reduced IgG2a levels in mice receiving the adjuvanted mRNA (**Figure 4E-G**). Cellular immunity was evaluated by quantifying antigen-reactive CD4^+^ and CD8^+^ T cells in the spleen. While the combinational adjuvants increased the percentage of antigen-reactive CD4^+^ T cells, anti-IFNAR treatment reduced this response to levels comparable to the non-adjuvant group (**Figure 4H, Supplemental Figure S4**), indicating that the enhanced CD4^+^ T cell response was largely dependent on type I IFN signaling. In contrast, although adjuvanted naked mRNA increased the percentage of CD8^+^ T cells compared to untreated controls, the adjuvants contributed minimally to CD8^+^ T cell responses (**Figure 4I**). Together, these findings demonstrate that type I IFN responses play a substantial role in the immunogenicity of adjuvanted naked mRNA vaccines, enhancing both innate and adaptive immune activation, consistent with the known ability of CpG to induce strong type I IFN responses (36–38, 49).

In order to obtain further mechanistic insights, we compared the gene expression profiles induced by Al-Phos and CpG with those induced by various adjuvants tested in **Figure 1**, which showed little or inhibitory effects on humoral or cellular immunity induction. In this experiment, we analyzed transcriptional profiles in the skin 24 hours after jet-injection of naked mRNA with or without these adjuvants. In quantitative PCR, we selected genes that are enriched in the combinational adjuvant group compared to the group without adjuvant in transcriptome analyses (**Figure 3**). Among the tested adjuvants, the combination of Al-Phos and CpG, as well as CpG alone, induced the highest or near-highest expression levels of type I IFN (*IFN-β*) and downstream genes of type I IFN signaling (*Mx1*, *Mx2*, *Isg15*, *STAT1*, *STAT2*, *Ifit2*) (**Figure 5**). The combinational adjuvants and CpG alone also enhanced expression of *Ifih1*, which encodes MDA5, an innate immune receptor positively regulated by type I IFN and other proinflammatory stimuli (50). These findings align with our previous results demonstrating the critical role of type I IFN signaling in naked mRNA jet-injection vaccination (**Figure 4**) and highlight the strong effects of CpG in inducing type I IFN, as previously reported (49).

**Figure 5.**
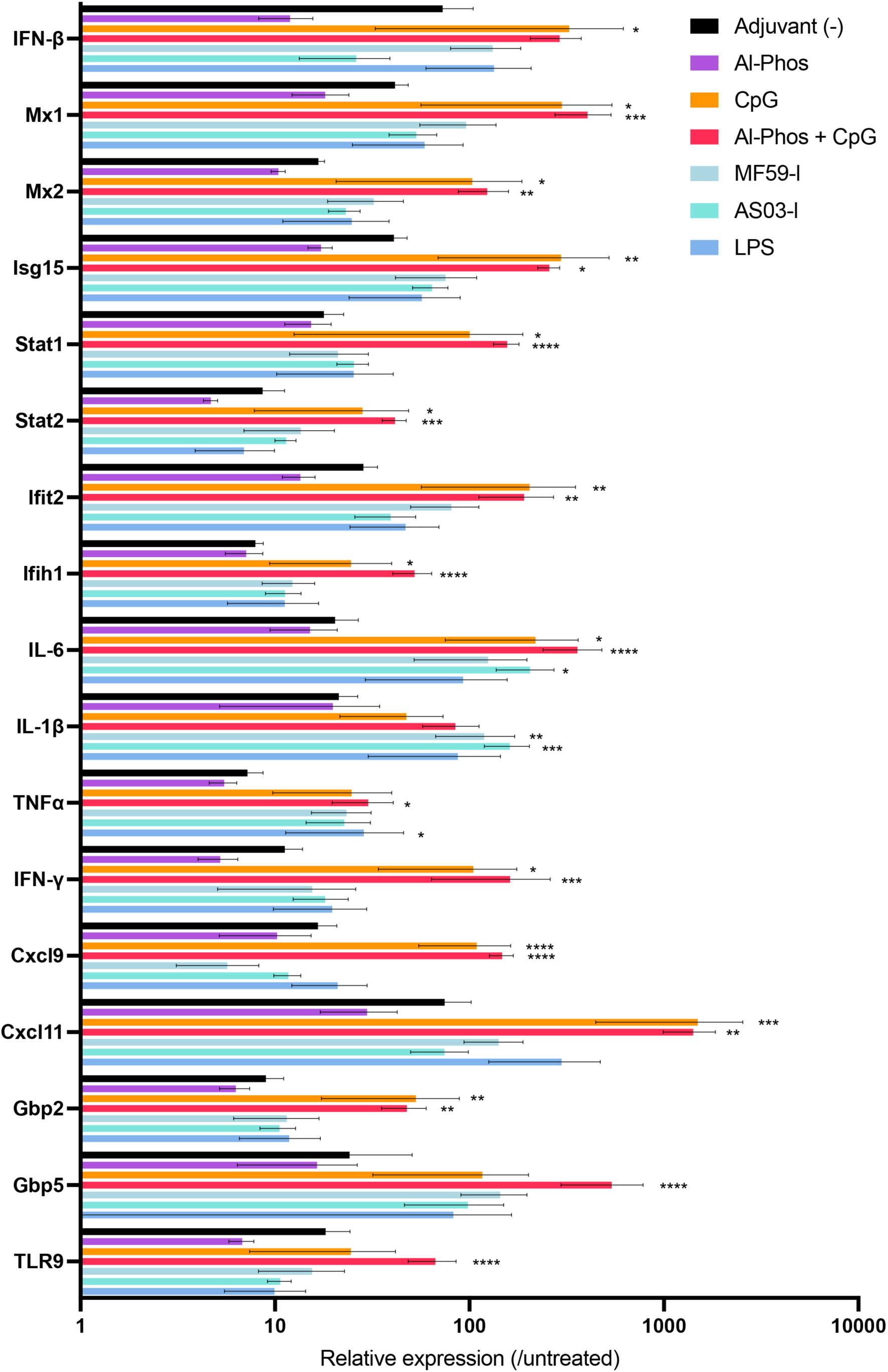
Gene expression profiles following vaccination with various adjuvants. Mice received naked mRNA with the indicated adjuvants or without adjuvant via jet-injection. At 24 hours post-immunization, the injected skin samples were collected for quantitative PCR. *n* = 4. Data are presented as mean ± s.d. Statistical differences compared to the control group without adjuvants were analyzed by using one-way ANOVA followed by Dunnett’s multiple comparisons test. *: *p* < 0.05; **: p < 0.01; ***: *p* < 0.001; ****: *p* < 0.0001.

Next, we evaluated the expression of additional cytokines (*IL-6*, *IL-1β, TNF-α, IFN-γ*), chemokines (*Cxcl9, Cxcl11*), inflammasome-related molecules primarily activated by IFN-γ (*Gbp2*, *Gbp5*), and an innate immune receptor (*TLR-9*) (**Figure 5**). The combinational adjuvants and CpG alone again induced the highest or near-highest expression levels of *IFN-γ*, the chemokines, the inflammasome-related molecules, and *TLR-*9, suggesting a possible contribution of these molecules to naked mRNA jet-injection vaccination. However, the effects on several cytokines, including *IL-6*, *IL-1β,* and *TNF-α*, were less pronounced across the tested adjuvants. Specifically, MF59-l, AS03-l, and LPS induced transcripts of these cytokines at levels comparable to those observed with the Al-Phos and CpG combination. Given MF59-l, AS03-l, and LPS failed to improve the vaccination efficacy of naked mRNA delivered by jet-injection, this finding suggests that these cytokines might provide limited benefit to the vaccination process, although further studies are needed to clarify and expand on their functional roles.

Jet-injection of naked mRNA without adjuvants increased transcript levels of the tested genes by several-fold to 100-fold compared with the untreated group (**Figure 5**). This finding is consistent with our previous report showing that jet-injection induced transient immune cell infiltration to the injected skin, presumably by providing physical stress (33). Meanwhile, the effect of Al-Phos alone on pro-inflammatory factor expression was masked by the pro-inflammatory effect of jet-injection, resulting in no further increase in transcript levels of the tested inflammatory genes, despite its substantial contribution to vaccination efficacy. These findings suggest that Al-Phos may function beyond simply inducing pro-inflammatory molecule expression during vaccination.

We also conducted transcript profiling in dLN (**Supplemental Figure S5**), which yielded results similar to those observed in the skin (**Figure 5**). The Al-Phos and CpG combination induced relatively high expression levels of type I IFN-related factors among the tested adjuvants. However, the superiority of the combinational adjuvant was less pronounced compared with that observed in the skin, probably due to the relatively modest enhancement of gene expression regardless of adjuvant type.

### Adjuvanted naked mRNA vaccine induces robust protection against SARS-CoV-2 challenge

The efficacy of a naked mRNA jet-injection vaccine supplemented with a combination of Al-Phos and CpG adjuvants was evaluated in a SARS-CoV-2 challenge model. Mice were vaccinated three times at two-week intervals, as repeated immunization enhanced antibody production efficiency (**Supplemental Figure S6A**). Of note, the beneficial effects of the combinational adjuvants remained evident even after three doses (**Supplemental Figure S6B**). From a safety perspective, body weights following three rounds of vaccination, with or without the combinational adjuvants, remained comparable to those of unvaccinated mice (**Supplemental Figure S6C**). Two weeks after the final vaccination, mice were intranasally challenged with SARS-CoV-2 (**Figure 6A**). To ensure effective infection, mice were sensitized by intranasally administration of an adenovirus expressing human ACE2 five days prior to the viral challenge (51, 52).

**Figure 6.**
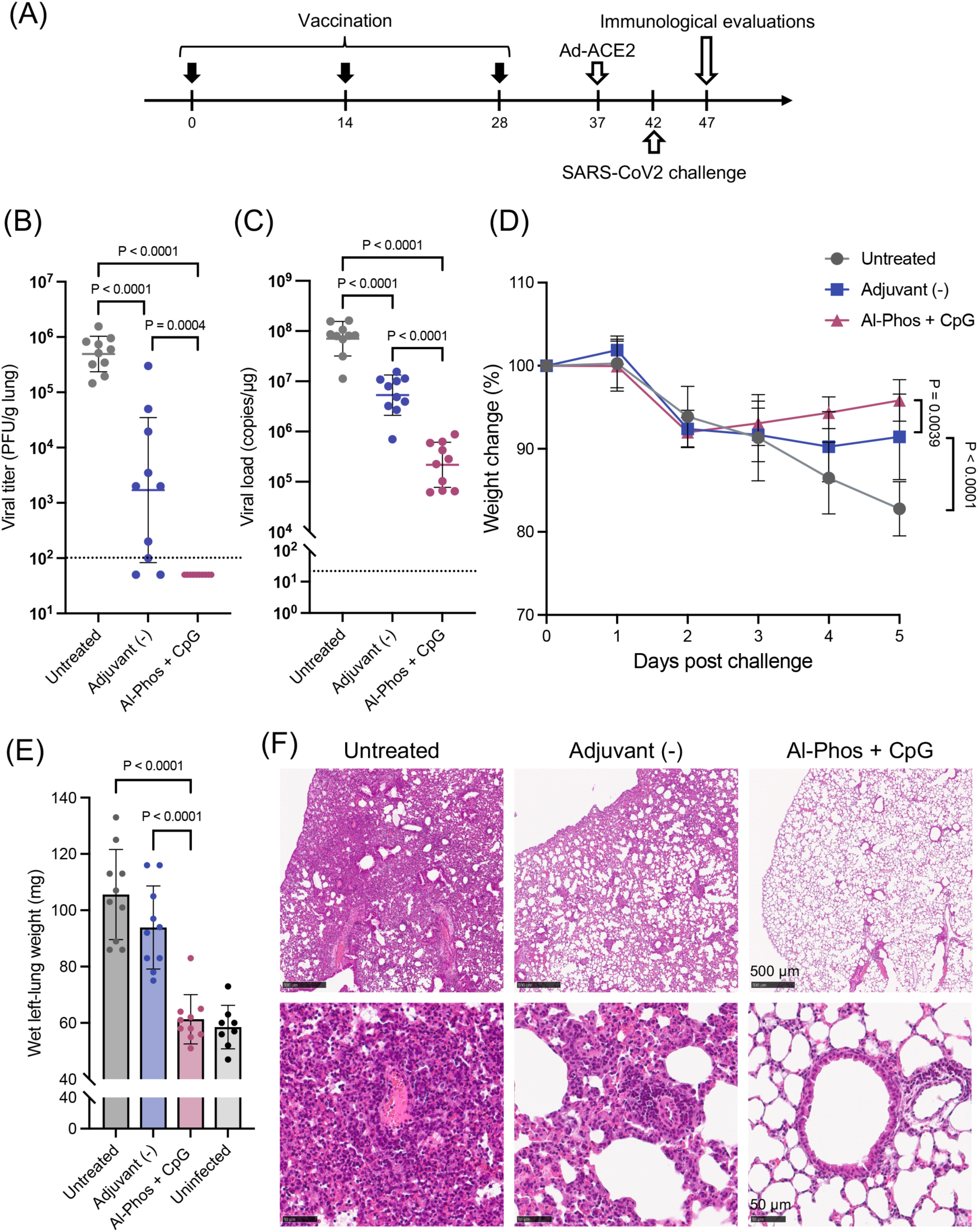
*Spike* mRNA vaccination against SARS-CoV-2 challenge. (**A**) Immunization and challenge schedule. Mice were immunized with naked *spike* mRNA with or without Al-Phos + CpG adjuvants via jet-injection, three times at 2-week intervals. Two weeks after the final dose, mice were intratracheally challenged with SARS-CoV-2. To permit infection, mice were sensitized 5 days before challenge by intratracheal administration of an adenovirus expressing human ACE2 (Ad-ACE2). Viral load measurement (**B, C**), lung weight measurement (**E**), and tissue section preparation (**F**) were conducted 5 days post-challenge. (**B, C**) SARS-CoV-2 viral burden in lung tissue quantified by plaque-forming assay (PFU) (**B**) and quantitative PCR of viral RNA (**C**). *n* = 10. Dashed lines indicate the limit of detection. (**D**) Body weight change following viral challenge. *n* = 10. (**E**) Wet weight of the left lung lobe. *n* = 10. (**F**) Representative H&E-stained lung sections post-challenge. Scale bars: 500 µm (upper), 50 µm (lower). Data are presented as geometric mean ± geometric s.d. for (**B, C**) and mean ± s.d. for (**D, E**). Statistical analyses were performed by using one-way ANOVA followed by Tukey’s multiple comparisons test.

In a plaque-forming assay conducted five days post-challenge, mice vaccinated with the adjuvanted naked mRNA showed viral loads in the lungs below the detection limit, demonstrating robust protective efficacy (**Figure 6B**). Although vaccination with unadjuvanted mRNA reduced viral load to some extent, substantial virus remained detectable at a significantly higher level compared to the adjuvanted naked mRNA (*p* = 0.0004 compared with adjuvanted naked mRNA; one-way ANOVA followed by Tukey’s test). Consistent with this result, PCR-based quantification revealed a ∼13-fold reduction in viral load following unadjuvanted mRNA vaccination and a 323-fold reduction with the adjuvanted formulation compared to the unvaccinated group. Thus, the use of the combinational adjuvants with naked mRNA significantly reduced viral load by 24-fold compared to naked mRNA without adjuvants [*p* < 0.0004 compared with Adjuvant (-); one-way ANOVA followed by Tukey’s test], highlighting their marked benefit (**Figure 6C**).

In addition to viral load, the general condition of the mice post-challenge was assessed by monitoring changes in body weights. Although all groups experienced weight loss within two days of viral challenge, mice vaccinated with the adjuvanted naked mRNA exhibited enhanced recovery compared to those receiving unadjuvanted mRNA or no vaccine (**Figure 6D**). Pulmonary edema was also evaluated by measuring wet lung weight, as viral infection can induce edema through tissue damage and inflammation. Lung weights in mice vaccinated with the adjuvanted mRNA were lower than those in unvaccinated or unadjuvanted groups and were comparable to those of uninfected mice. This finding suggests that vaccination mitigated lung edema (**Figure 6E**).

Consistently, histopathological examination further supported the protective effect of the combinational adjuvants (**Figure 6F**). Lungs from unvaccinated mice showed diffuse and severe lymphohistiocytic inflammatory infiltration with occasional neutrophils and reduced air content. Although unadjuvanted mRNA vaccination mitigated lung inflammation, moderate lymphohistiocytic infiltration was still observed in most regions of the lungs. These observations are consistent with our previous report demonstrating the efficacy of naked mRNA jet-injection in mitigating SARS-CoV-2 pneumonia in mice (33). In contrast, lungs from mice vaccinated with adjuvanted naked mRNA exhibited minimal histopathological abnormalities, with only mild lymphocyte-predominant inflammatory infiltration. These findings demonstrate that the adjuvanted naked mRNA vaccine not only greatly reduces viral load but also mitigates pathological changes following SARS-CoV-2 challenge. Collectively, the combination of Al-Phos and CpG substantially enhances the protective efficacy of naked mRNA jet-injection vaccines against SARS-CoV-2.

### Adjuvanted naked mRNA vaccine is effective against influenza virus

The versatility of adjuvanted naked mRNA vaccines was evaluated by targeting influenza A virus (A/California/7/2009 H1N1), as influenza viruses remain a major concern for ongoing or future epidemics including regional or global pandemics. To target hemagglutinin (HA), a widely used antigen in influenza vaccines, mRNA vaccine design offers two options: an endogenous membrane-bound form and a secreted form (**Figure 7A**) (53). The endogenous HA protein includes a secretion signal at the N-terminus and a transmembrane domain near the C-terminus. To create a secreted version, the transmembrane domain was replaced with a trimerization domain (i.e., an autonomous folding unit or foldon), which is commonly used in vaccines to enhance antigenicity. We first optimized antigen design by vaccinating mice with adjuvanted naked mRNA via jet-injection, administered three times at a two-week interval. Two weeks after the last vaccination, mRNA encoding the membrane-bound form induced stronger cellular immune responses in ELISpot assays compared to the secreted form, while both forms elicited comparable antibody levels (**Figure 7B, C**). Based on these results, we selected the membrane-bound form for subsequent analyses.

**Figure 7.**
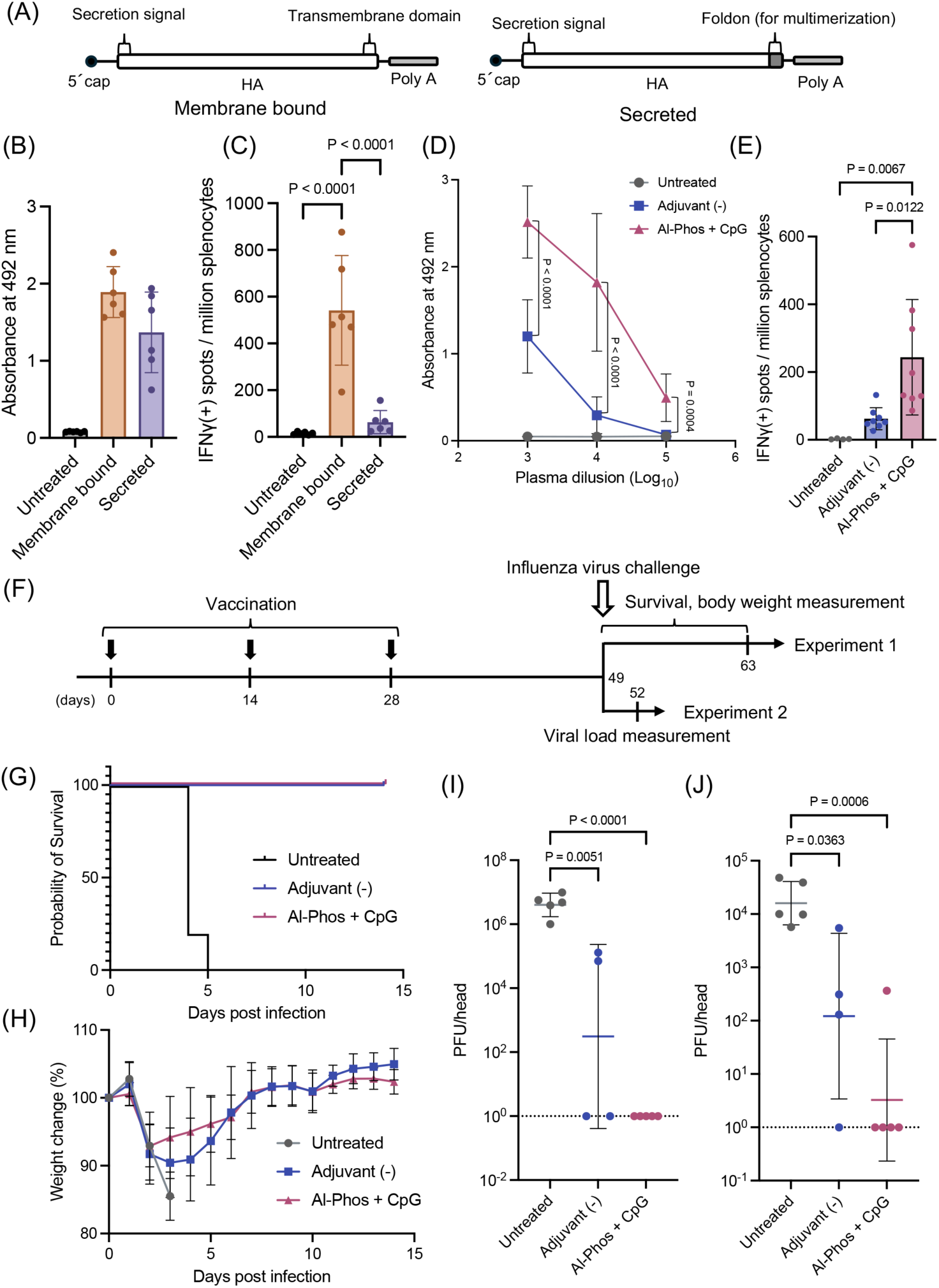
Hemagglutinin mRNA vaccination against influenza virus challenge. (**A-C**) Optimization of antigen design encoded by mRNA. (**A**) Schematic of mRNA constructs encoding membrane-bound or secreted hemagglutinin (HA) antigen. (**B, C**) Mice were immunized with naked mRNA encoding membrane-bound or secreted HA antigen with Al-Phos and CpG adjuvants via jet-injection three times at 2-week intervals. Blood and spleens were collected two weeks after the third dose. (**B**) Plasma HA-specific total IgG levels (absorbance values of plasma diluted 10,000-fold). *n* = 5. (**C**) IFN-γ-producing splenocytes by ELISpot. *n* = 5. (**D, E**) Mice were immunized with naked mRNA encoding membrane-bound HA antigen with or without Al-Phos and CpG adjuvants via jet-injection three times at 2-week intervals. Immune responses were evaluated two weeks after the third dose. (**D**) Dilution curves of HA-specific total IgG in plasma. *n* = 6. (**E**) IFN-γ-producing splenocytes by ELISpot. *n* = 5. (**F-J**) Viral challenge experiments. Mice were immunized 3 times at 2-week intervals and challenged with influenza virus three weeks after the third dose. (**F**) Experimental schedule. (**G, H**) Experiment 1: (**G**) Survival curves. *n* = 5. (**H**) Body weight change. *n* = 5. (**I, J**) Experiment 2: Influenza viral burden measured by plaque-forming assay (PFU) in nasal wash (**I**) and lung wash (**J**) [*n* = 5 for untreated and Al-Phos + CpG, *n* = 4 for Adjuvant (-)]. Dashed lines indicate the limit of detection. Data are presented as mean ± s.d. for (**B-E, H**) and geometric mean ± geometric s.d. for (**I, J**). Statistical analyses were performed by using one-way ANOVA followed by Tukey’s multiple-comparisons test.

Next, we evaluated the effect of adjuvant supplementation with Al-Phos and CpG in the naked mRNA jet-injection vaccine. As observed with *spike* mRNA, the combinational adjuvants enhanced both anti-HA antibody levels and the number of IFN-γ–secreting splenocytes responsive to HA (**Figure 7D, E**).

Finally, vaccine efficacy was assessed through a lethal influenza virus challenge model. Mice were vaccinated three times at two-week intervals, followed by viral challenge three weeks after the final dose (**Figure 7F**). All unvaccinated mice succumbed within five days (**Figure 7G**). In contrast, all vaccinated mice, regardless of adjuvant supplementation, survived the 14-day observation period and fully recovered their body weight (**Figure 7H**), demonstrating high efficacy of the naked mRNA jet-injection vaccine in this model. Consistently, plaque-forming assays conducted three days post-challenge revealed that both unadjuvanted and adjuvanted naked mRNA markedly reduced viral loads in nasal and lung washes compared to unvaccinated controls (**Figure 7I, J**). Meanwhile, adjuvanted mice tended to have lower viral loads compared to unadjuvanted mice. Among five mice receiving adjuvanted mRNA, virus was undetected in the lungs of all mice and detected in the nasal wash of only one mouse. In contrast, virus was detected in nasal or lung washes in half or more of the mice in the group without adjuvants. Collectively, these results demonstrate that supplementation with combinational adjuvants is a versatile strategy that enhances the efficacy of naked mRNA jet-injection vaccines, showing effectiveness against two distinct viral pathogens with well-documented potential for causing severe human diseases at an individual and population scale.

### Adjuvanted naked mRNA vaccines show safety benefits over LNP-based vaccines

To contextualize the efficacy of naked mRNA jet-injection vaccine, we compared it with a standard D-Lin-MC3-DMA–based LNP formulation by using the widely studied ovalbumin (OVA) antigen. LNPs were administered via intramuscular injection, the standard protocol in animal studies, clinical trials, and real-world applications (1–4). Mice were vaccinated twice at three-week intervals, and immunological assays were performed five weeks after the initial vaccination. Although LNPs induced higher levels of total anti-OVA IgG and IgG1 compared to naked mRNA supplemented with combinational adjuvants, both groups showed nearly comparable levels of IgG2a (**Figure 8A–F**). The balanced induction of IgG1 and IgG2a by the adjuvanted naked mRNA may reflect a favorable immune profile for reducing the risk of VAED (43, 44), although further studies are needed to confirm this. Additionally, both the adjuvanted naked mRNA vaccine and the LNP formulation elicited comparable cellular immune responses, as measured by ELISpot assays for IFN-γ–secreting splenocytes (**Figure 8G**).

**Figure 8.**
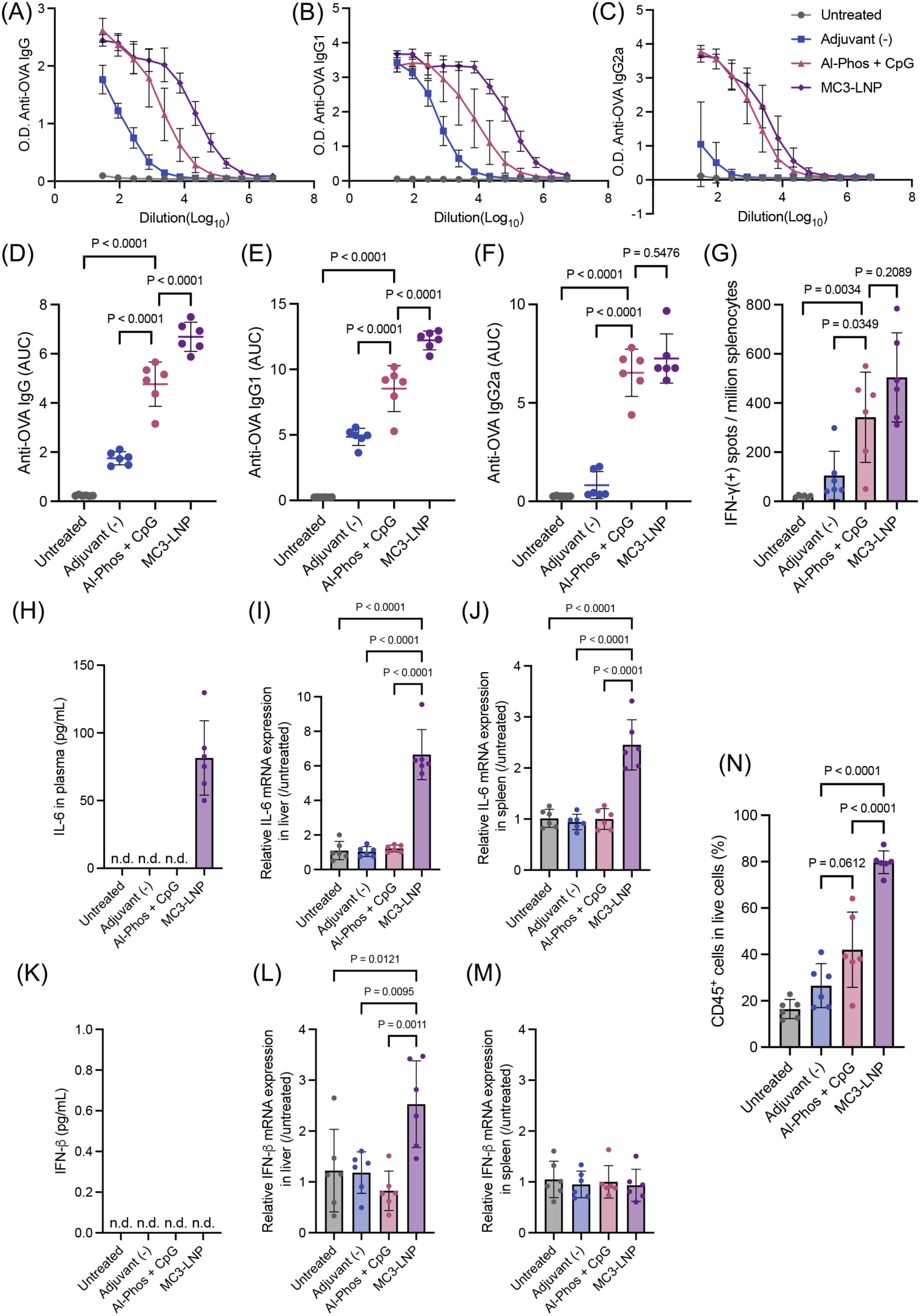
Vaccination efficacy and biosafety of adjuvanted naked mRNA and LNP. (**A–G**) Vaccination efficacy. Mice were immunized with *OVA* mRNA delivered either as naked mRNA with or without Al-Phos and CpG adjuvants via jet-injection or as mRNA formulated with D-Lin-MC3-DMA-based LNP (MC3-LNP) via intramuscular needle injection, twice at a 3-week interval. Vaccination efficacy was evaluated two weeks after the second dose. (**A–C**) Serial dilution curves of OVA-specific IgG (**A**), IgG1 (**B**), and IgG2a (**C**) in plasma determined by ELISA. *n* = 6. (**D–F**) Area under the curve (AUC) of the corresponding dilution curves in (**A–C**). IgG (**D**), IgG1 (**E**), IgG2a (**F**). *n* = 6. (**G**) IFN-γ–producing splenocytes quantified by ELISpot. *n* = 6. (**H–M**) Systemic biosafety analyses performed at 6 hours after naked mRNA jet-injection or intramuscular MC3-LNP needle injection. (**H, K**) Plasma levels of IL-6 (**H**) and IFN-β (**K**) quantified by ELISA. (**I, J, L, M**) Transcript levels of *IL-6* (**I, J**) and *IFN-β* (**L, M**) in the liver (**I, L**) and spleen (**J, M**) quantified by qPCR. *n* = 6. (**N**) Local safety analyses performed 6 hours after naked mRNA jet-injection or intradermal LNP needle and syringe injection. The percentage of CD45^+^ cells at the injection site was quantified by flow cytometry. Data are presented as mean ± s.d. Statistical analyses were performed by using one-way ANOVA followed by Tukey’s multiple comparisons test.

For safety analyses, we first assessed systemic inflammation by quantifying IL-6 and IFN-β protein levels in plasma, as well as the corresponding transcripts in the liver and spleen (**Figure 8H–M**). LNP injection increased plasma IL-6 protein levels and *IL-6* mRNA expression in both the liver and spleen (**Figure 8H–J**). These findings align with previous reports of systemic proinflammatory and toxicological responses at an mRNA dose of 10 μg in LNP formulations (54–56). In contrast, jet-injection of adjuvanted mRNA did not alter IL-6 protein or mRNA levels in plasma or systemic organs, although it did elevate IL-6 expression locally at the injection site and in dLNs (**Figures 3–5**). Notably, systemically released IL-6 contributes to vaccine-induced fever by upregulating cyclooxygenase-2 and prostaglandin E2, playing a key role in systemic reactogenicity (57). Evaluation of IFN-β also revealed the safety advantage of adjuvanted naked mRNA over LNP (**Figure 8K–M**). LNP increased IFN-β transcripts in the liver compared to the untreated control, while adjuvanted naked mRNA did not (**Figure 8L**). Meanwhile, in all vaccinated groups, IFN-β protein was undetected in plasma, and IFN-β transcript levels were comparable to those of untreated controls (**Figure 8K, M**).

Finally, we evaluated local reactogenicity by quantifying CD45^⁺^ leukocyte infiltration into dermal tissue following naked mRNA jet-injection or intradermal needle-and-syringe injection of LNP. Naked mRNA jet-injection, with or without adjuvants, showed only a modest increase in leukocyte counts (**Figure 8N**). In contrast, LNP injection resulted in greater leukocyte infiltration into the skin compared to naked mRNA jet-injection with or without combinational adjuvants. These results highlight the relatively low local reactogenicity of adjuvanted naked mRNA vaccines.

## Discussion

Naked mRNA-based vaccines offer a promising strategy to mitigate the relatively strong reactogenicity associated with LNP-based mRNA vaccines. To achieve sufficient efficacy, this approach must incorporate key LNP-like functionalities, including efficient mRNA delivery to APCs and strong adjuvanticity. While our previous study successfully enhanced mRNA delivery to APC-rich skin by using a jet injector (33), the present study focused on adjuvanticity and included a systematic screening to identify adjuvants that improve the efficacy of naked mRNA jet-injection vaccines without compromising safety profiles. As a result, the combination of Al-Phos and CpG markedly increased antibody production accompanied by enhanced germinal center formation and concurrently boosted CD4^+^ T cell responses (**Figure 2, 4H**). While naked mRNA jet-injection without adjuvants primarily induced IgG1, with minimal IgG2a production, supplementation with the combinational adjuvants led to a balanced induction of IgG1 and IgG2a. This balance may help reduce the risk of vaccine-associated enhanced disease (VAED) in certain infectious disease settings (43, 44). Furthermore, IgG2a plays a critical role in viral clearance due to its strong complement fixation ability and involvement in antibody-dependent cellular cytotoxicity (58).

Importantly, the adjuvanted naked mRNA vaccine induced IgG2a and cellular immunity at levels comparable to those induced by LNPs (**Figure 8A–G**). Although total IgG levels were lower than those induced by LNPs, the 10 μg LNP dose used in mice is considered excessive and was associated with systemic cytokine responses in our study (**Figure 8H–M**), as well as severe toxicity in other reports (54–56). In contrast, the adjuvanted naked mRNA vaccine did not induce detectable systemic inflammatory responses at the same dose, thereby highlighting its safety advantage. Although the combinational adjuvants enhanced local pro-inflammatory responses, local reactogenicity remained milder compared to that with LNP vaccines (**Figure 8N**).

From a practical standpoint, the adjuvanted naked mRNA vaccine demonstrated effectiveness against both coronaviruses and influenza viruses (**Figures 6, 7**), two major concerns for future respiratory-transmitted pandemics. Notably, vaccination reduced viral loads in the lungs to background levels in plaque-forming assays for both viruses, indicating sufficient immunization to inhibit viral pneumonia. For clinical translation, it is worth noting that both Al-Phos and CpG are clinically licensed, although the CpG design used in our system differs from that in approved vaccines (36–38). Moreover, the jet injector used in our study is adaptable to various species, including humans, by simply adjusting the injection force (42, 59). Finally, it has already been tested in a clinical trial for a naked DNA vaccine, showing only modest local reactogenicity (34). As an mRNA vaccine platform, jet-injection of naked mRNA induced comparable antibody production efficiency in both mice and non-human primates, demonstrating its potential for application in large animal species (33).

Notably, prior investigations on SARS-CoV-2 and COVID-19 vaccination have introduced targeted vaccine design concepts with streamlined deployment (60, 61), identified conserved epitopes to support broad protective coverage (62), and developed quantitative and modeling-based approaches to understand viral load dynamics, vaccine performance, and pandemic trajectories (63–67). These complementary studies provide a conceptual framework for the present work, in which rational adjuvant and delivery design enhances the immunogenicity and safety profile of mRNA-based vaccination, thereby supporting the further translational development of adaptable vaccine platforms for future infectious disease threats.

Regarding the mechanisms of action, the Al-Phos and CpG combination enhanced the expression of type I IFN-related genes in the injected skin and dLN (**Figure 3**). Blocking type I IFN signaling diminished the ability of the combinational adjuvants to enhance DC activation, IgG2a induction and CD4^+^ T cell responses (**Figure 4**). These findings highlight the essential role of type I IFN signaling in driving robust innate and adaptive immune responses elicited by the combinational adjuvants. Notably, CpG alone was sufficient to induce transcripts related to type I IFN signaling at levels comparable to those induced by the combinational adjuvants (**Figure 5**), although it failed to improve cellular immunity induction efficiency (**Figure 1D**). These findings suggest that Al-Phos plays a role beyond enhancing type I IFN responses. Importantly, one should note that Al-Phos alone did not increase the transcript levels of any tested immune-related genes compared to the control group injected with unadjuvanted mRNA, suggesting that Al-Phos contributes to vaccination process independently of inducing pro-inflammatory factor expression. Aluminum adjuvants are well known for their antigen depot effect, adsorbing antigens to enable slow release or enhanced delivery to dLN (36–38, 68). Although negatively charged Al-Phos may not adsorb negatively charged mRNA, it could capture antigen proteins expressed from mRNA or other immune-related molecules, thereby enhancing their bioavailability. Additional functions of aluminum adjuvants, such as the release of damage-associated molecular patterns (DAMPs) (69), may likely contribute to improving the efficacy of naked mRNA jet-injection vaccines as well.

Another relevant finding of this study is that clinically approved adjuvants or their derivatives, which are known to enhance the efficacy of inactivated, toxoid, or recombinant protein vaccines (36–38, 70) failed to enhance humoral or cellular immune responses in naked mRNA jet-injection vaccines (**Figure 1C–E**). The route of delivery may be a potential factor explaining this discrepancy, as adjuvant functions are typically evaluated after intramuscular injection, whereas this study used the intradermal route. However, intradermal injection of these adjuvants successfully increased the expression of proinflammatory molecules in this study (**Figure 5**, **Supplemental Figure S5**). Other studies have also reported pro-inflammatory effects of these adjuvants following intradermal delivery (71). Therefore, the use of mRNA itself, rather than differences in administration routes, may be a more plausible explanation for the limited or inhibitory effects of adjuvants observed in this proof-of-concept translational study.

Previous studies have revealed negative impacts of excessive proinflammatory responses on mRNA vaccine efficacy. One plausible mechanism is the suppression of antigen protein expression from mRNA due to inflammation (30, 40, 41). However, this is unlikely to be the main cause of the limited or negative effects of adjuvants in this study, as mixing adjuvants had minimal impact on mRNA delivery efficiency in the skin except for MF59-l (**Figure 1A, B**). Proinflammatory responses may also impair the immunization process after antigen expression, for example by suppressing T cell functions. This issue has been studied primarily in the context of type I IFN signaling, which can either enhance or diminish mRNA vaccine efficacy depending on vaccination conditions (7, 20, 41, 45–47, 72–74). This dual role may be explained by the context of immunostimulation, such as the kinetics of type I IFN signaling and antigen presentation. Specifically, antigen presentation preceding type I IFN signaling is preferred for optimal T cell function. These contexts of immunostimulation may differ between adjuvant types and between conventional and mRNA vaccines, perhaps explaining the complex functions of adjuvants observed in this study.

Despite reports of negative impacts from excessive type I IFN responses in LNP-based mRNA vaccines reported (7, 41, 74), this study demonstrated a clear benefit of type I IFN induction by adjuvants in naked mRNA jet-injection vaccines. While the precise mechanisms underlying this discrepancy remain unclear, it is reasonable to assume that the overall intensity of the pro-inflammatory response to vaccines could influence its function. Within this speculative context, the relatively low pro-inflammatory response elicited by naked mRNA jet-injection compared to LNPs (**Figure 8**) might explain the positive effect of type I IFN observed in this study. Although this study revealed the essential role of type I IFN signaling in enhancing IgG2a production and CD4^+^ T cell responses elicited by the adjuvanted naked mRNA vaccines, blockade of type I IFN signaling did not affect total IgG or IgG1 levels (**Figure 4E-G**), suggesting the involvement of additional signaling pathways. In this regard, the combination of Al-Phos and CpG increased the expression of pro-inflammatory factors beyond type I IFN-related molecules, including *IFN-γ*, chemokines, inflammasome-related molecules, and *TLR-*9 (**Figure 5**). These factors might also contribute to the enhanced induction of antigen-specific antibodies.

In conclusion, we successfully improved the efficacy of naked mRNA vaccines by combining jet-injection for efficient skin delivery with an optimal adjuvant combination of Al-Phos and CpG. The adjuvanted naked mRNA vaccines provided robust protection against SARS-CoV-2 and influenza A virus. Gratifyingly, this strategy induced no detectable systemic reactogenicity and only modest local reactogenicity, ensuring a favorable safety profile. Moreover, the use of clinically licensed adjuvants and jet injectors with proven applicability in humans and large animals highlights the clinical translatability of our strategy. Finally, this proof-of-concept translational study also provides valuable mechanistic insights into adjuvant selection for mRNA vaccines, thereby revealing the critical role of type I IFN in mRNA vaccines and the differences in optimal adjuvants between conventional and mRNA-based platforms. Given the high likelihood of future global pandemics such the 1918 influenza pandemic or the recent COVID-19, this technology and the findings reported here will serve as a translational framework for development into human patient applications.

## Methods

Detailed methods are described in **Supplemental Methods**.

### Sex as a biological variable

Sex is not considered as a biological variable in this study. Only female mice were used.

### Immunization

Female BALB/c mice (6–8 weeks old) were intradermally injected in the left flank with 10 µg of N1-methyl-pseudouridine-modified mRNA encoding *spike*, *HA*, or *OVA*, in a total volume of 20 µL by using a jet injector (Actranza® lab.; Daicel Corporation, Tokyo, Japan), as previously described (33). For adjuvanted formulations, mRNA was mixed with Adju-Phos® (final concentration 0.1% w/w; Al-Phos), ODN 1585 VacciGrade™ (final concentration 1 mg/mL; CpG), MPLA VacciGrade™ (final concentration 0.5 mg/mL; LPS), AddaVax (mixed 1:1 with the mRNA solution; MF59-l), AddaS03 (mixed 1:1 with the mRNA solution; AS03-l), or a combination of Adju-Phos® (final concentration 0.1% w/w) and ODN 1585 VacciGrade™ (final concentration 0.5 mg/mL). For blocking type-I IFN or IL-6 receptor, anti-IFNAR-1 antibody (clone MAR1-5A3, Bio X Cell) or anti-mouse IL-6R antibody (clone 15A7, Bio X Cell) was injected intraperitoneally at 500 µg per mouse one day before immunization.

### Evaluation of humoral and cellular immune responses

Enzyme-linked immunosorbent assay (ELISA) was performed to quantify antibodies. Briefly, 96-well plates were coated with OVA protein (20 μg/mL, Sigma-Aldrich), spike protein (1 μg/mL, Sino Biological), or influenza A H1N1 (A/California/07/2009) hemagglutinin / HA protein (His tag) (2 μg/mL, Sino Biological, 11085-V08H). Antigens were incubated with diluted plasma for 2 hours at room temperature (RT), followed by incubation with HRP-conjugated goat anti-mouse IgG (1:8000, HAF018, R&D Systems), IgG1 (1:10000, ab97240, Abcam), or IgG2a (1:10000, ab97245, Abcam) antibodies for one hour at RT. Cellular immunity was assessed by using mouse IFN-γ ELISpot PLUS kits (Mabtech) and following the manufacturer’s instructions. In the protocol, splenocytes (5 × 10^5^ cells/well) were stimulated with 0.25 μg/mL OVA peptide pool (Miltenyi Biotec), 2 μg/mL spike peptide pool (JPT Peptide Technologies), or 2 μg/mL Influenza A, Hemagglutinin, California (H1N1), PepMix (JPT Peptide Technologies) for 18–24 hours at 37 °C.

### RNA sequencing (RNA-seq)

Skin tissues were finely minced with sterile scissors, homogenized in 900 μL TRIzol reagent using a PowerMasher II homogenizer (NIP) for 5 min, and processed with the RNeasy Plus Universal Mini Kit (QIAGEN). dLN samples were homogenized in 500 μL RLT buffer containing 1% β-mercaptoethanol with a Micro Smash (MS-100, TOMY) at 3,000 rpm for 60 sec, and total RNA was extracted by using the RNeasy Mini Kit (QIAGEN). RNA integrity was verified with an Agilent 4200 TapeStation system; all samples had RIN values ≥ 7.0 (range: 7.0–8.8). Library preparation and sequencing were performed by Macrogen Japan Corp. Libraries were constructed with the TruSeq Stranded mRNA Library Prep Kit (Illumina) and sequenced on a NovaSeq X platform (101-bp paired-end). The sequencing provider performed quality control, read trimming, alignment to the mm10 mouse reference genome, and transcript quantification. TPM values were used for expression analysis. All samples achieved Q30 scores >97% and mapping rates >97%. Differential expression analysis was conducted by using the DESeq2, comparing adjuvanted (Al-Phos + CpG) versus non-adjuvanted groups. Genes with |fold change| ≥ 2 and false-discovery rate (FDR) < 0.05 (Benjamini–Hochberg correction) were considered significantly differentially expressed. Pathway analysis was performed by using GSEA with clusterProfiler (v4.14.6), ranking genes by Wald statistic. Mouse Hallmark gene sets from MSigDB were obtained via msigdbr (v25.1.1), and 100,000 permutations were used for significance testing.

### Challenge with SARS-CoV-2

Challenge experiments with SARS-CoV-2 were performed as previously described (33). Briefly, five days after intranasal inoculation of mice with 5 × 10^7^ FFU/animal of rAd5-hACE2 (51, 52), mice were infected with 10^5^ PFU/50 µL/animal of an early circulating SARS-CoV-2 strain (TY/WK-521/2020; GISAID ID: EPI_ISL_408667). For plaque assays, lung homogenates were centrifuged at 3,000 × g for 10 min at 4 °C. Vero E6/TMPRSS2 cells were incubated with the supernatants at 37 °C for 1 hour. After washing, cells were overlaid with DMEM containing 10% FBS and 0.6% agarose and incubated for 48 hours at 37 °C. Cells were then fixed and stained with 1% crystal violet. For quantitative PCR, the amount of N gene in lung homogenates was quantified.

### Challenge with influenza virus

The A/California/7/2009 (H1N1)pdm09 (X-179A) virus was propagated in 10-day-old embryonated chicken eggs. Mice were intranasally challenged with 10 LD_50_ (50 μL per mouse) of the virus via the left nostril. Virus titers in nasal and lung wash samples were determined by plaque assay, as previously described (75). Briefly, serially diluted nasal and lung wash samples were inoculated onto Madin-Darby Canine Kidney (MDCK) cell monolayers. After one hour, wells were washed twice with PBS and overlaid with 2 mL of agar medium containing acetylated trypsin from bovine pancreas (Sigma-Aldrich, St. Louis, MO, USA) at 10 μg/mL. After two days, cells were fixed and stained with crystal violet. These animal experiments were conducted in biosafety level two animal facilities.

### Statistics

One-way or two-way analysis of variance (ANOVA) followed by Tukey’s or Dunnett’s multiple comparisons test was performed for comparisons among three or more groups by using Graph Pad Prism 10.6.1. Unpaired Student’s t-tests were performed for comparisons between two groups. Details of statistical methods are provided in the corresponding figure legends, as indicated.

### Study approval

All animal experiments were conducted in strict compliance with animal husbandry and welfare regulations under ethical approval from the Innovation Center of NanoMedicine (iCONM), Kawasaki Institute of Industrial Promotion (approval number: A20-010-6), Institute of Science Tokyo (approval number: A2025-114C2), Tokyo Metropolitan Institute of Medical Science (approval number: 27-071), and the National Institute of Infectious Diseases (approval number: 124112), all in Japan.

## Data availability

All data are available upon reasonable request. RNA sequencing data have been deposited in NCBI’s Gene Expression Omnibus under accession number GSE314153.

## Acknowledgments

This work was supported in part by the Leading Advanced Projects for Medical Innovation (JP21gm0010008 to SU), Research Program on Emerging and Re-emerging Infectious Diseases (JP21fk0108620 to SU), and the Strategic Center of Biomedical Advanced Vaccine Research and Development for Preparedness and Response (SCARDA) (223fa827006h0001 to SU) from Japan Agency for Medical Research and Development (AMED), the Open Innovation Platform for Industry-Academia Cocreation (COI-NEXT) Program (Grant Number: JPMJPF2022 to SU) from the Japan Science and Technology Agency (JST), Grants-in-Aid for Challenging Research (Pioneering) (23K1748 to SU) and Scientific Research (A) (21H04962 to SU) from the Ministry of Education, Culture, Sports, Science and Technology, Japan (MEXT), the Program for Promotion of Fundamental Studies in COVID-19 of the Tokyo Metropolitan Government (to FY, KK, MK, and SU), Nanken-Kyoten, Science Tokyo (to SU), and Medical Research Center Initiative for High Depth Omics, Science Tokyo (to SU), and by the Levy-Longenbaugh Donor-advised Fund (to RP and WA). We thank Erika Mochizuki and Reiko Shiratori (Institute of Science Tokyo) for their technical assistance and Dr. Helen Pickersgill (Life Science Editors) for professional editing of the manuscript.

## Author Contributions

NQ, SU designed research; NQ, JI, FY, KS, KM, AH, AFA, LH, ZX performed research; NQ, JI, KM, TS, RP, WA, KK, HH, MK, SU provided intellectual insights or reagents; NQ, JI, FY, KS, AH, RP, WA, KK, SU analyzed data; and NQ, JI, FY, KS, RP, WA, KK, SU wrote and/or edited the manuscript.

