## Supplemental Data for "Combinational adjuvants delivered by ink-jet potentiate naked mRNA vaccines for robust protection against infectious diseases"

**Contents**

Supplemental Methods …………………………………………………………………………….. 2

Supplemental Figures …………………………………………………………………………….. 8

Supplemental Table …………………………………………………………………………….. 14

Supplemental References …………………………………………………………………………….. 15

### Supplemental Methods

#### Materials

The Actranza® lab. jet injector was purchased from Daicel Corporation (Tokyo, Japan). *Firefly luciferase* mRNA (*fLuc* mRNA) was obtained from Trilink BioTechnologies (San Diego, CA, USA). N1-methyl-pseudouridine (m1Ψ)-modified *Ovalbumin (OVA)* mRNA, m1Ψ-modified *SARS-CoV-2 spike* mRNA, and m1Ψ-modified *IFNα11* mRNA were synthesized by GenScript (Piscataway, NJ, USA). Al-Phos (Adju-Phos®, vac-phos-250), CpG (ODN 1585 VacciGrade™, vac-1585-1), MF59-l (AddaVax, vac-adx-10), AS03-l (AddaS03, vac-as03-10), and LPS (MPLA VacciGrade™, vac-mpls,) were purchased from Invivogen (San Diego, CA, USA). The Luciferase Assay System was purchased from Promega Co. (Fitchburg, WI, USA). OVA protein was purchased from Sigma-Aldrich (St. Louis, MO, USA). SARS-CoV-2 spike protein was purchased from Sino Biological Inc. (Beijing, China). PepTivator Ovalbumin epitope mix was purchased from Miltenyi Biotec (Nordrhein-Westfalen, Germany). PepMix™ SARS-CoV-2 (S) was purchased from JPT Peptide Technologies (Berlin, Germany). Anti-IFN-γ ELISpot PLUS kits were purchased from Mabtech (Nacka Strand, Sweden). Horseradish peroxidase (HRP)-conjugated goat anti-mouse IgG antibody was purchased from R&D systems (Minneapolis, MN, USA). HRP-conjugated goat anti-mouse IgG1 and IgG2a antibodies were purchased from Abcam (Cambridge, UK). *InVivo*MAb anti-mouse IFNAR-1 and InVivoMAb anti-mouse IL-6R antibodies were purchased from Bio X Cell (Lebanon, NH, USA). Uncoated IL-6 and IFN-β ELISA kits were purchased from R&D Systems (Minneapolis, MN, USA).

#### *In vivo* mRNA delivery

Female BALB/c mice (6–8 weeks old) were purchased from Sankyo Labo Service Corporation (Tokyo, Japan). Mice were intradermally injected in the left flank with 3 μg of *fLuc* mRNA for bioluminescence imaging or 10 μg of mRNA encoding *spike*, *HA*, or *OVA* for immunization, in a total volume of 20 μL using a jet injector under isoflurane anesthesia. For adjuvanted formulations, mRNA was mixed with Adju-Phos® (final concentration 0.1% w/w; Al-Phos), ODN 1585 VacciGrade™ (final concentration 1 mg/mL; CpG), MPLA VacciGrade™ (final concentration 0.5 mg/mL; LPS), AddaVax (mixed 1:1 with the mRNA solution; MF59-l), AddaS03 (mixed 1:1 with the mRNA solution; AS03-l), or a combination of Adju-Phos® (final concentration 0.1% w/w) and ODN 1585 VacciGrade™ (final concentration 0.5 mg/mL). For bioluminescence imaging, mice were intraperitoneally injected with 200 μL of 15 mg/mL VivoGlo Luciferin (Promega), followed by bioluminescence imaging using an in vivo imaging system (IVIS, PerkinElmer). Lipid nanoparticles (LNPs) were prepared from D-Lin-MC3-DMA, DSPC, cholesterol, and PEG2000-DMG as previously described (1) and intramuscularly or intradermally injected into mice with needle. For blocking of type-I interferon or IL-6 receptor, anti-IFNAR-1 antibody (clone MAR1-5A3, Bio X Cell) or anti-mouse IL-6R antibody (clone 15A7, Bio X Cell) was injected intraperitoneally at 500 μg per mouse in 200 μL PBS one day before immunization.

#### Enzyme-linked immunosorbent assay (ELISA) for antibody quantification

Clear Flat-Bottom Immuno Nonsterile 96-Well Plates (Thermo Fisher) were coated with OVA protein (20 μg/mL, Sigma-Aldrich), spike protein (1 μg/mL, Sino Biological), or influenza A H1N1 (A/California/07/2009) hemagglutinin / HA protein (His tag) (2 μg/mL, Sino Biological, 11085-V08H) in carbonate buffer (pH 9.6) overnight at 4 °C. Plates were washed with PBS containing 0.05% Tween 20 and blocked with 1% BSA in PBS for 1 hour at room temperature. After three washes, 50 µL of plasma serially diluted three-fold in diluent buffer (1% BSA, 0.05% Tween 20 in PBS) was added and incubated for 2 hours at room temperature. Plates were then washed three times and incubated with 50 µL of HRP-conjugated goat anti-mouse IgG (1:8000, HAF018, R&D Systems), IgG1 (1:10000, ab97240, Abcam), or IgG2a (1:10000, ab97245, Abcam) antibodies in diluent buffer for 1 hour at room temperature. For **Figures 2** and **8** and **Supplementary Figure S6**, plates were then washed three times, and treated with 100 µL of 3,3′,5,5′-tetramethylbenzidine (TMB) substrate. The reaction was stopped by adding 50 µL of 2 N sulfuric acid, and absorbance was measured at 450 nm using a SPARK 20M plate reader (Tecan Trading AG, Männedorf, Switzerland). For **Figures 1** and **7** and **Supplementary Figure S1**, after washing, 50 µL of o-phenylenediamine dihydrochloride (OPD) substrate solution was added. The reaction was stopped by adding 12.5 µL of 3 M hydrochloric acid, and optical density (OD) was measured at 492 nm using a SPARK 20M plate reader.

**Cell isolation from spleen**

Spleens were collected from immunized mice, mechanically dissociated into single-cell suspensions, and passed through 40 µm cell strainers. Suspensions were layered over Lympholyte-M (Cedarlane Labs) and centrifuged at 800 × g for 30 min at room temperature to isolate lymphocytes. The interface layer was harvested, washed twice with PBS, and resuspended in complete RPMI-1640 medium supplemented with 1 mM sodium pyruvate, 10 mM HEPES, and 50 µM β-mercaptoethanol. Cells were counted using a Countess 3 Automated Cell Counter (Thermo Fisher) and diluted to the desired concentration for subsequent assays.

**IFN-γ enzyme-linked immunosorbent spot (ELISpot) assay**

IFN-γ ELISpot assays were performed using mouse IFN-γ ELISpot PLUS kits (Mabtech) according to the manufacturer's instructions. Precoated plates were washed five times with PBS and blocked with complete medium for 30 min at room temperature. Splenocytes ($5 \times 10^5$ cells/well) were plated and stimulated with 0.25 µg/mL OVA peptide pool (Miltenyi Biotec), 2 µg/mL spike peptide pool (JPT Peptide Technologies), or 2 µg/mL Influenza A, Hemagglutinin, California (H1N1), PepMix (JPT Peptide Technologies) for 18–24 hours at 37 °C. After incubation, cells were removed by washing five times with PBS, and biotinylated anti-mouse IFN-γ detection antibody (1:1000 dilution in PBS containing 0.5% FBS) was added for 2-hour incubation at room temperature. Plates were washed again and incubated with streptavidin-HRP (1:1000 in PBS containing 0.05% FBS) for 1 hour. Following five additional washes with PBS, 100 µL per well of TMB substrate solution was added and incubated in the dark until distinct spots appeared. The reaction was stopped by rinsing with deionized water, and plates were dried overnight. Spots were counted using an ELISpot reader (AID GmbH, Strassberg, Germany).

**Analysis of GC B cells and Tfh cells in draining lymph nodes**

Draining lymph nodes (dLNs) were mechanically dissociated by gently pressing tissues through 40-µm nylon strainers using syringe plungers to obtain single-cell suspensions. Cells were washed with PBS and stained for viability using Fixable Aqua Dead Cell Stain (1:1000 in PBS, L34957, Invitrogen) for 30 min at 4 °C. After washing with MACS buffer (PBS with 2% FBS and 2 mM EDTA), Fc receptors were blocked with anti-CD16/32 antibody

(1:50 dilution; clone 93, BioLegend) for 10 min at 4 °C. Surface staining was performed in MACS buffer for 40 min at 4 °C using the following antibodies: Brilliant Violet 605 anti-mouse CD3 (clone 17A2, BioLegend), PerCP/Cy5.5 anti-mouse CD4 (clone GK1.5, BioLegend), APC/Cy7 anti-mouse CD279 (PD-1; clone 29F.1A12, BioLegend), FITC anti-mouse CD185 (CXCR5; clone L38D7, BioLegend), PE anti-mouse/human CD45R/B220 (clone RA3-6B2, BioLegend), and Brilliant Violet 421 anti-mouse CD38 (clone 90, BioLegend). Cells were washed twice with MACS buffer and fixed with IC Fixation Buffer (eBioscience) for 20 min at room temperature. Data were acquired using a CytoFLEX LX flow cytometer (Beckman Coulter) and analyzed with FlowJo software.

**RNA sequencing (RNA-seq)**

Skin tissues were finely minced with sterile scissors, homogenized in 900 μL TRIzol reagent using a PowerMasher II homogenizer (NIP) for 5 min, and processed with the RNeasy Plus Universal Mini Kit (QIAGEN). dLN samples were homogenized in 500 μL RLT buffer containing 1% β-mercaptoethanol using a Micro Smash (MS-100, TOMY) at 3,000 rpm for 60 sec, and total RNA was extracted using the RNeasy Mini Kit (QIAGEN). RNA integrity was verified using an Agilent 4200 TapeStation system; all samples had RIN values ≥ 7.0 (range: 7.0–8.8).

Library preparation and sequencing were performed by Macrogen Japan Corp. Libraries were constructed using the TruSeq Stranded mRNA Library Prep Kit (Illumina) and sequenced on a NovaSeq X platform (101-bp paired-end). The sequencing provider performed quality control, read trimming, alignment to the mm10 mouse reference genome, and transcript quantification. TPM values were used for expression analysis. All samples achieved Q30 scores >97% and mapping rates >97%.

Differential expression analysis was conducted using DESeq2, comparing adjuvanted (Al-Phos + CpG) versus non-adjuvanted groups. Genes with |fold change| ≥ 2 and FDR < 0.05 (Benjamini–Hochberg correction) were considered significantly differentially expressed. Pathway analysis was performed using GSEA with clusterProfiler (v4.14.6), ranking genes by Wald statistic. Mouse Hallmark gene sets from MSigDB were obtained via msigdbr (v25.1.1), and 100,000 permutations were used for significance testing.

Data visualization was performed in R (v4.4.1). The top 15 enriched pathways were displayed as bar plots showing normalized enrichment scores (NES) colored by −log10(FDR) (**Figure 3A, B**). Overlay plots of running enrichment scores were generated to compare four key cytokine signaling pathways: IFN-α, IFN-γ, IL6/JAK-STAT3, and TNF-α/NF-κB (**Figure 3C, D**). Volcano plots were annotated with 12 curated immune pathways from MSigDB Mouse Hallmark and GO Biological Process collections. Gene labels were shown for representative DEGs belonging to the 12 immune pathways meeting either $-\log_{10}$(FDR) in the top 10% among all DEGs, or |log2FC| > 2.5 (**Figure 3E, F**). Expression levels of the top 50 DEGs were grouped by pathway and displayed as bar plots of log2(TPM + 1) values with standard deviation ($n$ = 3) (**Figure 3G, H**). Figures were generated using ggplot2 (v4.0.0) and enrichplot (v1.26.6).

RNA sequencing data have been deposited in the NCBI's Gene Expression Omnibus under GEO accession number of GSE314153.

**Measurement of cytokine mRNA levels by quantitative PCR**

Total RNA from dLNs and injection-site skin tissues was extracted as described above. Complementary DNA (cDNA) was synthesized using the ReverTra Ace qPCR RT Master Mix Kit (TOYOBO). Relative gene expression levels of proinflammatory cytokines were determined in dLNs and skin samples by quantitative PCR (qPCR) using SYBR Green Universal Master Mix (Thermo Fisher) on a QuantStudio Real-Time PCR System (Applied Biosystems). Primer sequences used for mRNA quantification are listed in **Supplementary Table S1**. Target gene expression was normalized to β-actin using the $2^{-\Delta\Delta Ct}$ method, and results were expressed as relative fold changes compared with control samples.

**Analysis of activation of DCs in draining lymph nodes**

Single-cell suspensions were stained with Fixable Aqua Dead Cell Stain (1:1000 in PBS, L34957, Invitrogen) for 30 min at 4 °C to assess viability. After washing, Fc receptors were blocked with anti-CD16/32 antibody (1:50 dilution; clone 93, BioLegend) for 10 min at 4 °C. Cells were stained in MACS buffer for 40 min at 4 °C with FITC anti-mouse CD45 (clone 30-F11, BioLegend), Brilliant Violet 650 anti-mouse CD11c (clone N418, BioLegend), PE anti-mouse I-A/I-E (clone M5/114.15.2, BioLegend), Brilliant Violet 421 anti-mouse CD40 (clone 3/23, BioLegend), and APC anti-mouse CD86 (clone GL-1, BioLegend) antibodies. After staining, cells were washed twice with MACS buffer and fixed with IC Fixation Buffer (eBioscience) for 20 min at room temperature. Flow cytometric data were acquired using a CytoFLEX LX flow cytometer (Beckman Coulter) and analyzed with FlowJo software.

**Flow cytometry analysis of cytokine-producing cells in spleens**

Splenocytes were plated at $2 \times 10^6$ cells per well in 96-well U-shaped plates and stimulated with a 2 μg/mL spike overlapping peptide pool (JPT Peptide Technologies) and 2.5 μg/mL anti-CD28 (Tonbo #40-0281-M001) for 6 hours in complete RPMI 1640 medium. After 1 hour of stimulation, GolgiPlug (BD Biosciences) was added to each well, and the cells were incubated for an additional 5 hours. Following restimulation, cells were collected, washed with PBS, and stained with Fixable Aqua Dead Cell Stain (1:1000 in PBS, L34957, Invitrogen) for 30 min at 4 °C to assess viability. After washing with FACS staining buffer (2% FBS in PBS), cells were blocked with anti-CD16/32 antibody (1:50 dilution; clone 93, BioLegend) for 10 min at 4 °C, then stained with APC anti-mouse CD3ε (clone 17A2, BioLegend), PE anti-mouse CD4 (clone GK1.5, eBioscience), FITC anti-mouse CD8a (clone 53-6.7, BioLegend) antibodies for 40 min at 4 °C for surface marker detection. Cells were fixed for 20 min at room temperature and permeabilized twice using the Fixation/Permeabilization buffer set (eBioscience) and stained intracellularly with APC-eFluor™ 780 anti-IFN-γ antibodies (clone XMG1.2, eBioscience) in permeabilization buffer for 60 min at room temperature. Flow cytometric data were acquired with a CytoFLEX LX flow cytometer (Beckman Coulter) and analyzed using FlowJo software.

**Challenge with SARS-CoV-2**

Challenge experiments with SARS-CoV-2 were performed as previously described (1). Briefly, five days after intranasal inoculation of mice with $5 \times 10^7$ FFU/animal of rAd5-hACE2 (2, 3), mice were infected with $1 \times 10^5$ PFU/50 μL/animal of an early circulating SARS-CoV-2 strain (TY/WK-521/2020; GISAID ID: EPI_ISL_408667). Lungs were collected at 5 days post-infection. For plaque assays, lung homogenates were centrifuged at 3,000 × g for 10 min at 4 °C. Vero E6/TMPRSS2 cells were incubated with the supernatants at 37 °C for 1 hour. After washing with DMEM, cells were overlaid with DMEM containing 10% FBS and 0.6% agarose and incubated for 48 hours at 37 °C. Cells were then fixed and stained with 1% crystal violet. For qPCR, the amount of N gene in lung homogenates was quantified using the following primers and probe:

- Forward primer: 5'-GACCCCAAAATCAGCGAAAT-3'
- Reverse primer: 5'-TCTGGTTACTGCCAGTTGAATCTG-3'
- Probe: 5'-(FAM)-ACCCCGCATTACGTTTGGTGGACC-(BHQ-1)-3'

For histopathology, 4-μm paraffin sections of lungs were prepared and stained with hematoxylin and eosin.

**Challenge with influenza virus**

The A/California/7/2009 (H1N1)pdm09 (X-179A) virus was propagated in 10-day-old embryonated chicken eggs. Mice were intranasally challenged with 10 $LD_{50}$ (50 μL per mouse) of the virus via the left nostril. Three days post-infection, half of the mice were euthanized under deep isoflurane anesthesia, and serum, nasal wash, and lung lavage samples were collected. Nasal washes were obtained by flushing the nasal cavity through a cannula inserted from the posterior nasal aperture with 1 mL of wash buffer (PBS containing 0.1% bovine serum albumin, 10 U/mL penicillin, and 10 μg/mL streptomycin). Lung lavage fluid was collected by performing three washes through a cannula inserted into the trachea using 2 mL of wash buffer per wash. The remaining mice were monitored for 14 days post-infection to assess body weight changes and survival. Mice that lost more than 20% of their initial body weight were euthanized.

Virus titers in nasal and lung wash samples were determined by plaque assay, as previously described (4). Briefly, Madin-Darby Canine Kidney (MDCK) cells (American Type Culture Collection; CCL-34) were maintained at 37°C in a humidified atmosphere with 5% $CO_2$ in Minimum Essential Medium (MEM; Life Technologies, Carlsbad, CA, USA) supplemented with 10% fetal bovine serum (Thermo Fisher Scientific, Grand Island, NY, USA) and penicillin-streptomycin (100 U/mL penicillin, 100 μg/mL streptomycin; Life Technologies). MDCK cells were seeded into 6-well plates (Corning) and grown to confluence. Serial 10-fold dilutions of nasal and lung wash samples were prepared, and 200 μL of each dilution was inoculated onto MDCK monolayers. After 1 hour at 37 °C in 5% $CO_2$, wells were washed twice with PBS and overlaid with 2 mL of agar medium containing acetylated trypsin from bovine pancreas (Sigma-Aldrich, St. Louis, MO, USA) at 10 μg/mL. After 2 days at 37 °C in 5% $CO_2$, cells were fixed and stained with crystal violet. Plaques were counted, and virus titers were expressed as plaque-forming units per mouse (PFU/head). These animal experiments were conducted in biosafety level two animal facilities.

**In vivo systemic inflammation and biosafety**

Plasma was separated from blood for cytokine quantification using Uncoated Mouse IL-6 and IFN-β ELISA kits (R&D Systems) according to the manufacturer's instructions. Briefly, 96-well plates were coated with 100 μL per well of diluted capture antibody overnight at 4 °C. After washing and blocking, standards and 50 μL of plasma samples were added to each well and incubated for 2 hours at room temperature. Wells were then washed and incubated sequentially with diluted detection antibody for 1 hour and streptavidin-HRP for 20 min at room temperature. After washing, TMB substrate solution was added and incubated in the dark for 20 min. The reaction was stopped with 2 N sulfuric acid, and absorbance was measured at 450 nm using a Spark plate reader (Tecan, Switzerland). For proinflammatory cytokines expressed in spleen and liver, gene expression levels were measured by qPCR as described above.

**Supplemental Figures**

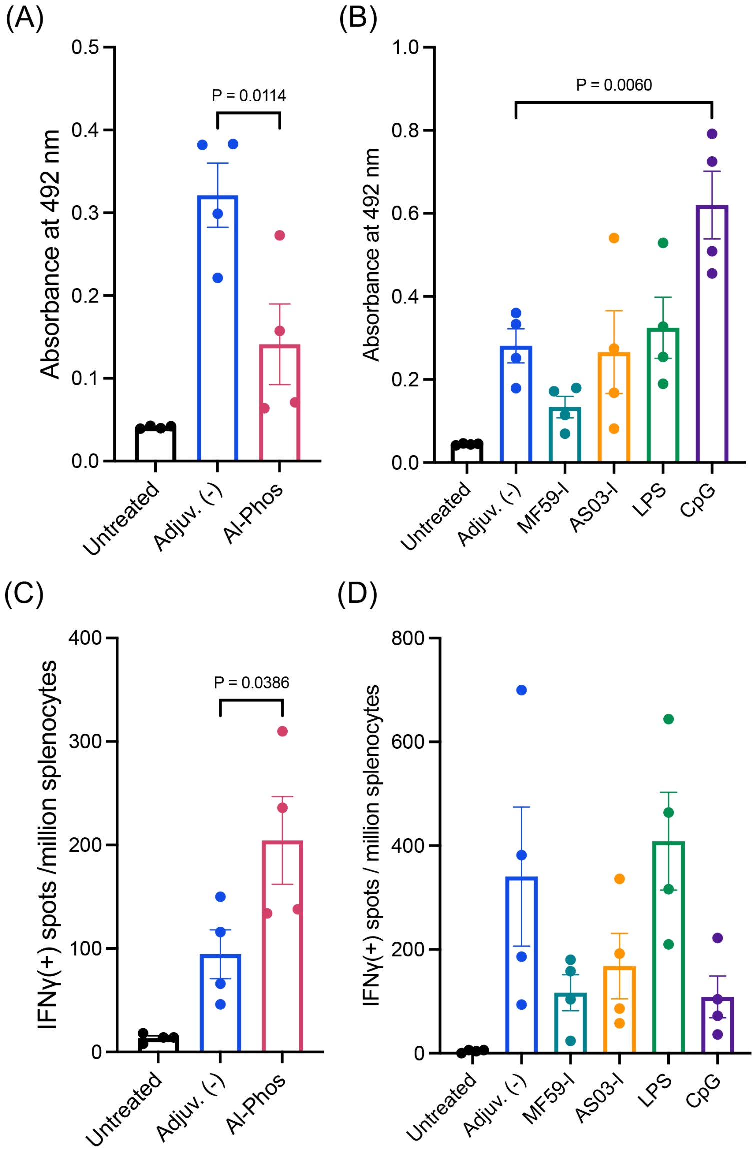


**Supplemental Figure S1. Influence of adjuvants on naked mRNA jet-injection vaccines.** Mice were immunized with naked *spike* mRNA with or without adjuvants via jet-injection, administered twice at a 3-week interval. Immune responses were evaluated two weeks after the second dose. (**A, B**) Spike-specific total IgG levels in plasma determined by ELISA. Absorbance values for plasma diluted 10,000-fold are shown. $n = 4$. (**C, D**) IFN-γ-producing splenocytes assessed by ELISpot. $n = 4$. Data are presented as mean ± s.e.m. Statistical analyses were performed by using one-way ANOVA followed by Dunnett's multiple comparisons test. These results were used to generate **Figure 1C-E**.

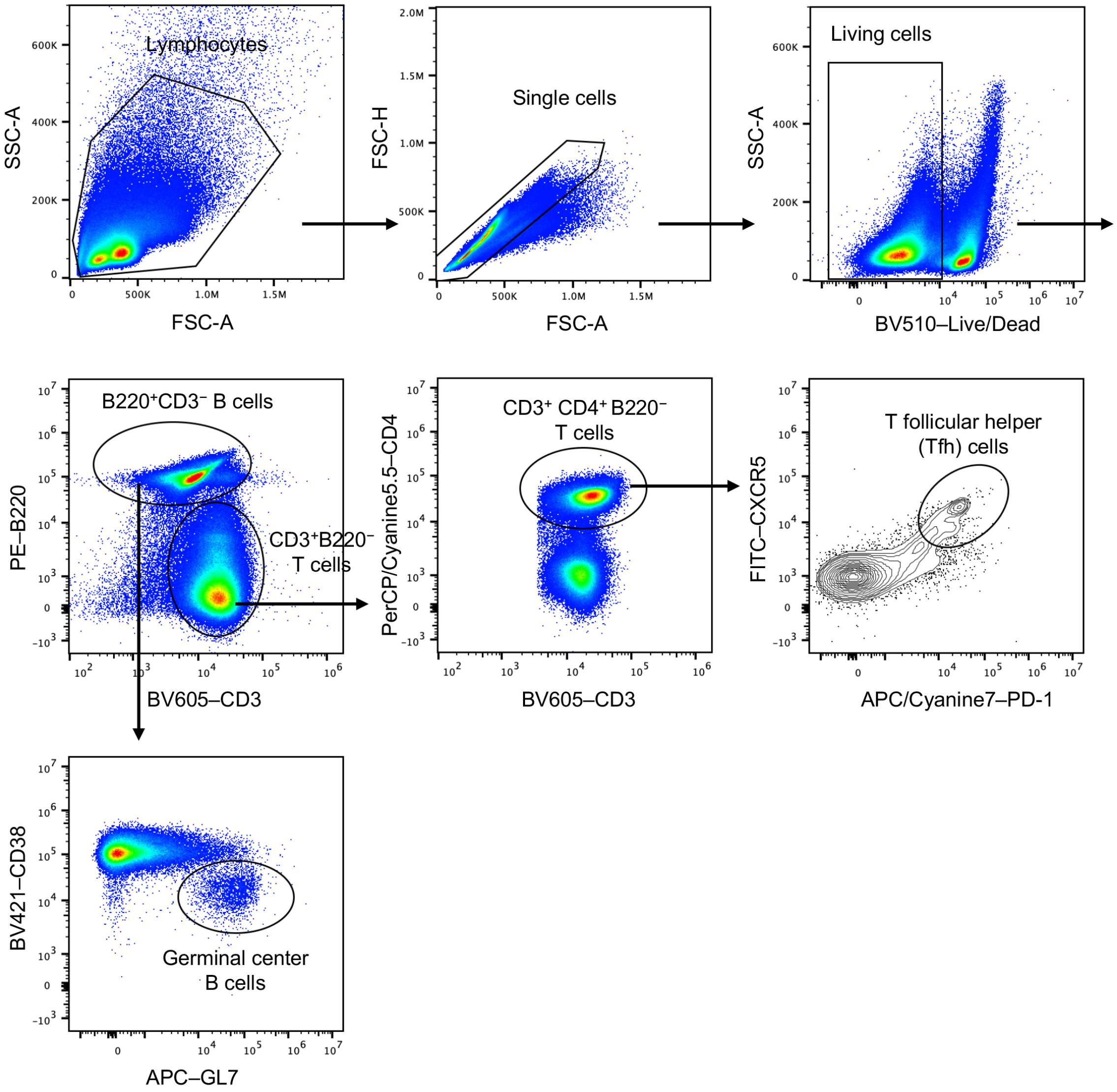


**Supplemental Figure S2. Gating strategy for flow cytometric analysis of Tfh cells and GC B cells in dLNs.** Representative gating plots are shown from a mouse on day 7 after immunization with the adjuvanted naked mRNA. Gates were applied to identify Tfh ($CD4^+CXCR5^+PD\text{-}1^+$) cells and GC B cells ($B220^+GL7^+CD38^-$).

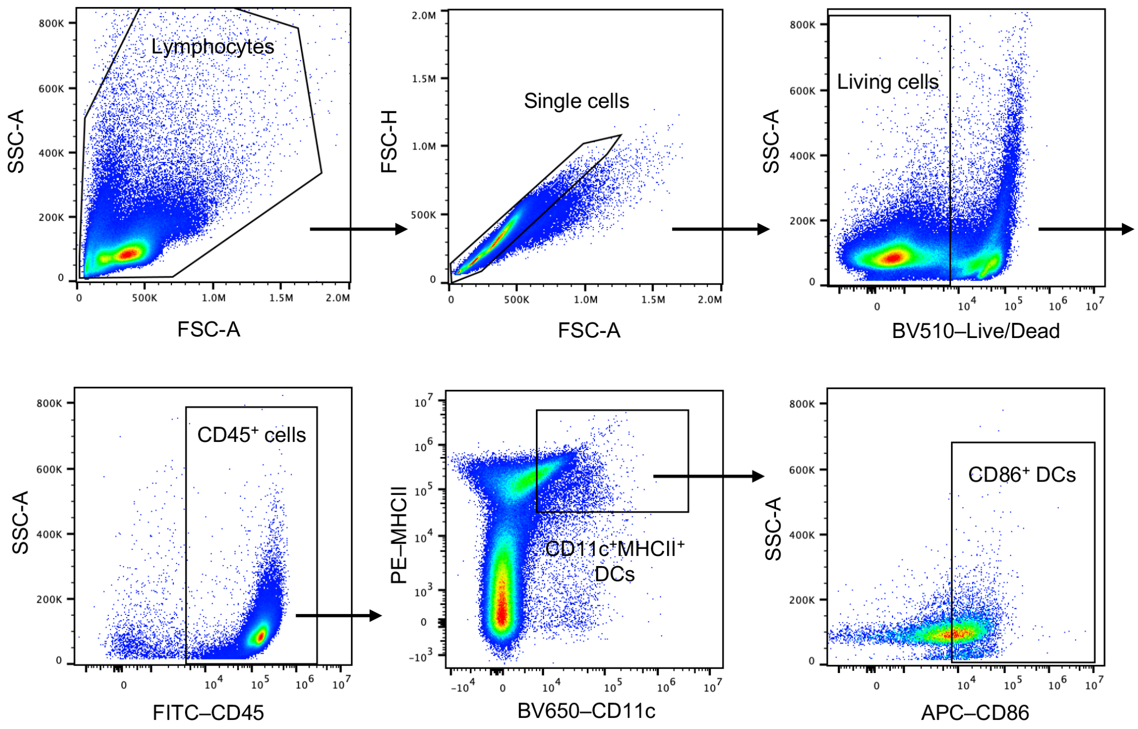


**Supplemental Figure S3. Gating strategy for flow cytometric analysis of DCs activation in dLNs.** Representative gating plots are shown from a mouse on day 1 after immunization with the adjuvanted naked mRNA.

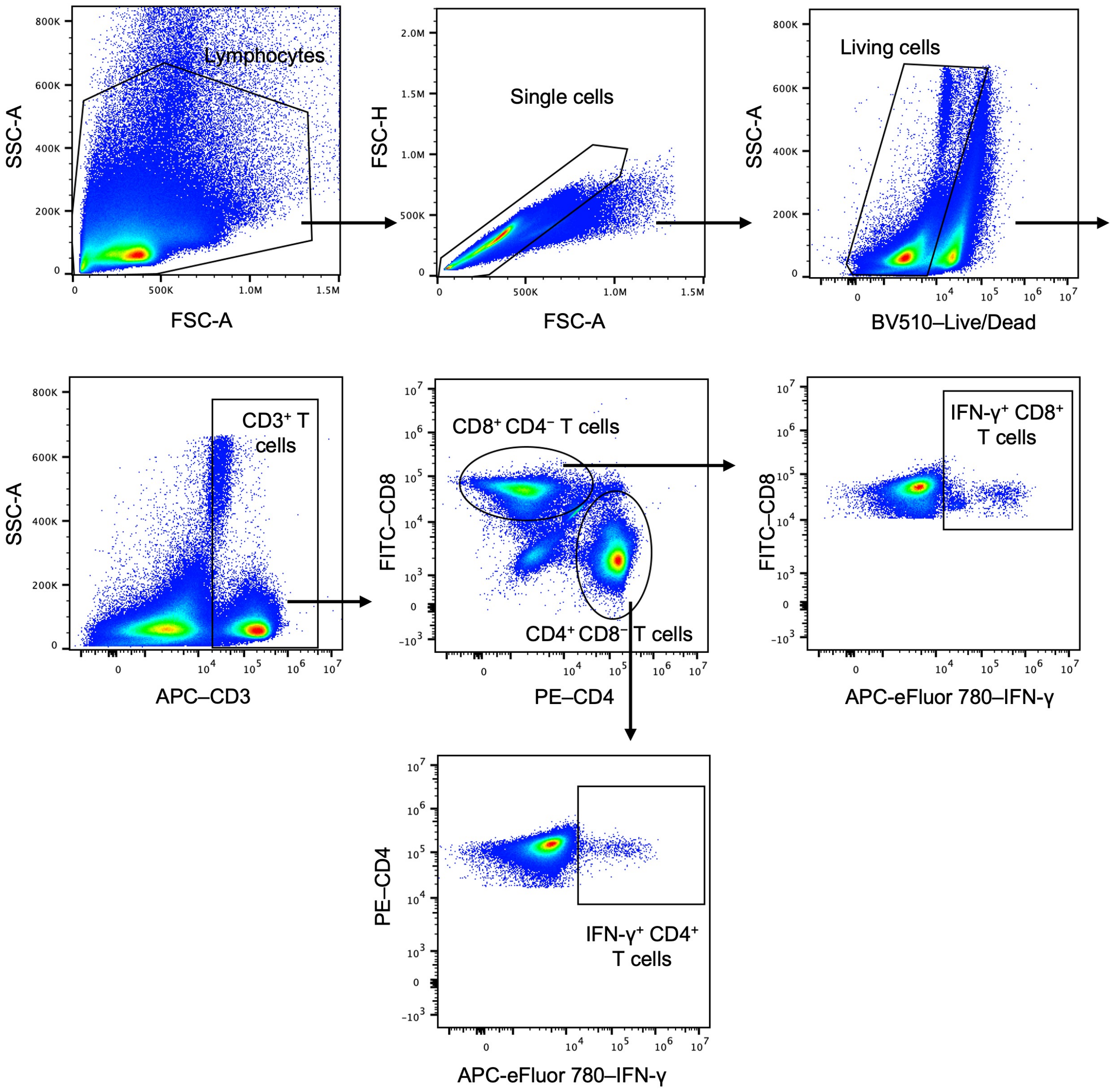


**Supplemental Figure S4. Gating strategy for flow cytometric analysis of IFN-γ–secreting splenocytes.** Representative gating plots are shown from a mouse 35 days after the first vaccination with adjuvanted naked mRNA.

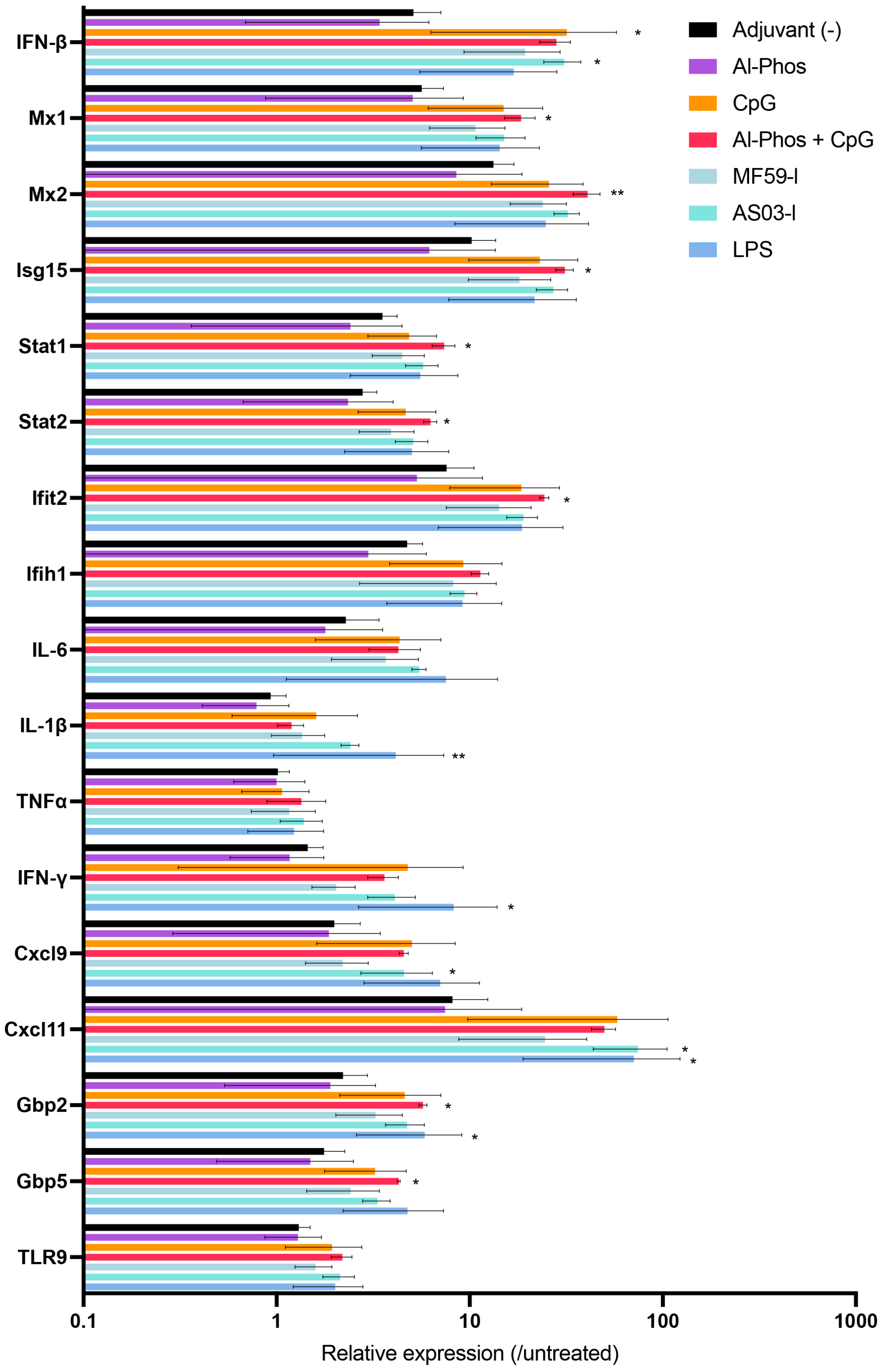


**Supplemental Figure 5. Gene expression profiles in dLNs following vaccination with various adjuvants.** Mice received naked mRNA with the indicated adjuvants or without adjuvant via jet-injection. At 24 hours post-immunization, dLNs were collected for quantitative PCR. $n = 4$. Data are presented as mean ± s.d. Statistical differences compared to the control group without adjuvants were analyzed by using one-way ANOVA followed by Dunnett's multiple comparisons test. *: $p < 0.05$; **: $p < 0.01$.

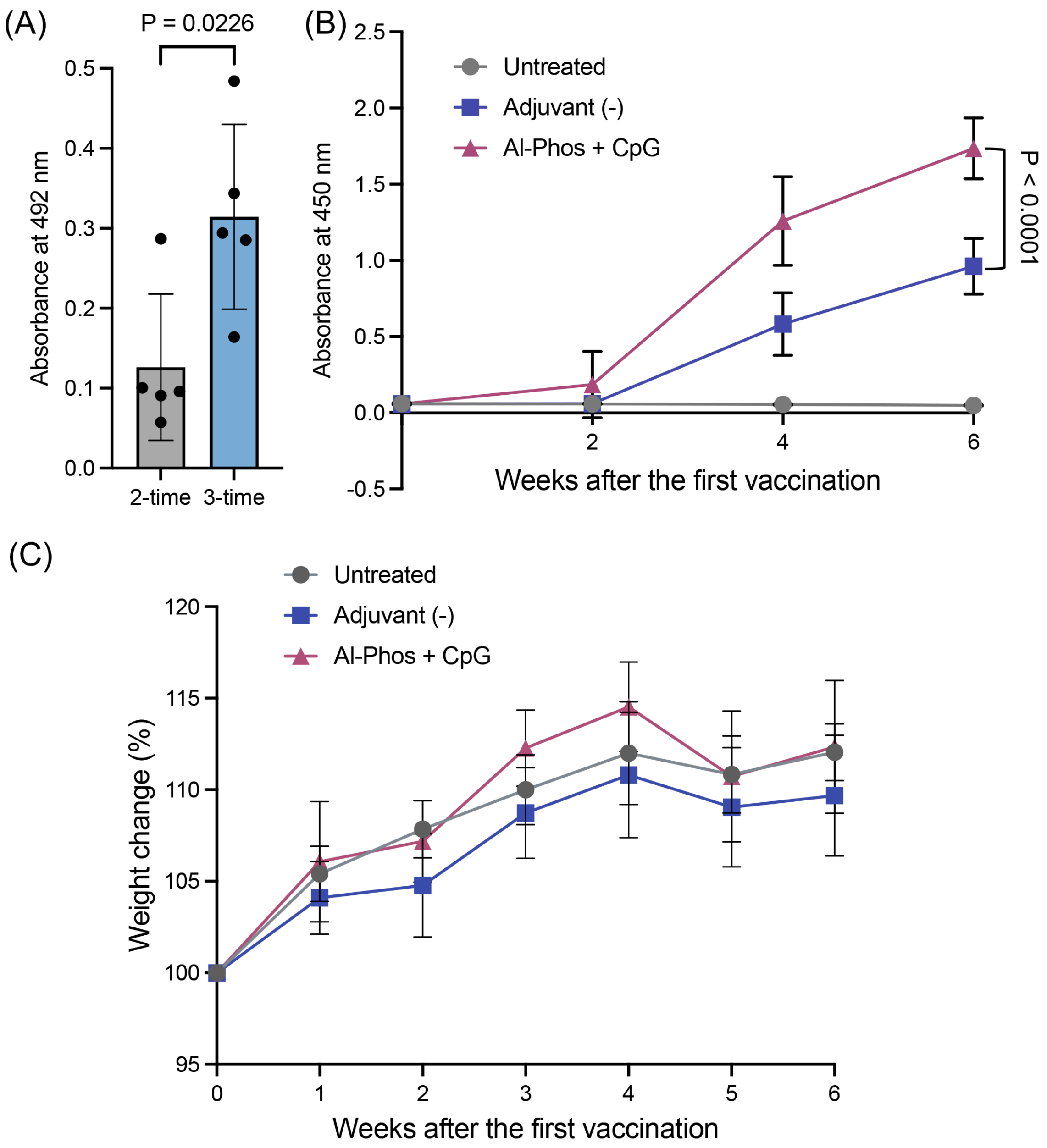


**Supplemental Figure S6. Frequent vaccination with naked mRNA jet-injection. (A)** Mice were immunized with naked *spike* mRNA without adjuvants either twice at a 3-week interval or three times at 2-week intervals. Spike-specific total IgG levels in plasma were assessed 6 weeks after the first dose and are presented as absorbance values at 10,000-fold dilution. $n = 5$. Data are represented as the mean ± s.d. Statistical analysis was performed by two-tailed unpaired Student's t-test. (**B, C**) Mice were immunized with naked *spike* mRNA with or without Al-Phos and CpG adjuvants via jet-injection, three times at a 2-week interval. (**B**) Spike-specific total IgG levels in plasma determined by ELISA at 0, 2, 4, and 6 weeks after the first dose, presented as absorbance values at 1,000-fold dilution. $n = 10$. (**C**) Body weight changes in mice vaccinated with mRNA or adjuvanted mRNA during the immunization schedule. $n = 10$. Data are represented as the mean ± s.d. Statistical analysis was performed by one way ANOVA followed by Tukey's multiple comparisons test.

**Supplemental Table**

**Supplemental Table 1 Primer sequences used in this study.**

| **Gene name** | **Forward primers** | **Reversed primers** |
|---|---|---|
| ACTB | CATTGCTGACAGGATGCAGAAGG | TGCTGGAAGGTGGACAGTGAGG |
| Ifit2 | AGTACAACGAGTAAGGAGTCACT | AGGCCAGTATGTTGCACATGG |
| Mx1 | GACCATAGGGGTCTTGACCAA | AGACTTGCTCTTTCTGAAAAGCC |
| Mx2 | GAGGCTCTTCAGAATGAGCAAA | CTCTGCGGTCAGTCTCTCT |
| Stat1 | GATCGCTTGCCCAACTCTTG | ACTGTGACATCCTTGGGCTG |
| Stat2 | TCCTGCCAATGGACGTTCG | GTCCCACTGGTTCAGTTGGT |
| Isg15 | GGTGTCCGTGACTAACTCCAT | TGGAAAGGGTAAGACCGTCCT |
| Cxcl11 | GCTTTCTCGATCTCTGCCAT | AACAGGAAGGTCACAGCCAT |
| Cxcl9 | GAGCAGTGTGGAGTTCGAGG | TCCGGATCTAGGCAGGTTTG |
| Gbp2 | CTGCACTATGTGACGGAGCTA | GAGTCCACACAAAGGTTGGAAA |
| Gbp5 | CAGACCTATTTGAACGCCAAAGA | TGCCTTGATTCTATCAGCCTCT |
| Tlr9 | ATGGTTCTCCGTCGAAGGACT | GAGGCTTCAGCTCACAGGG |
| Ifih1 | GCCTGGAACGTAGACGACAT | TGGTTGGGCCACTTCCATTT |
| IL-6 | TACCACTTCACAAGTCGGAGGC | CTGCAAGTGCATCATCGTTGTTC |
| IFN-β | GCCTTTGCCATCCAAGAGATGC | ACACTGTCTGCTGGTGGAGTTC |
| TNFα | GGTGCCTATGTCTCAGCCTCTT | GCCATAGAACTGATGAGAGGGAG |
| IFN-γ | ATGAACGCTACACACTGCATC | CCATCCTTTTGCCAGTTCCTC |
| IL-1β | TGGACCTTCCAGGATGAGGACA | GTTCATCTCGGAGCCTGTAGTG |
